# Impact of in vivo protein folding probability on local fitness landscapes

**DOI:** 10.1101/590398

**Authors:** Matthew S. Faber, Emily E. Wrenbeck, Laura R. Azouz, Paul J. Steiner, Timothy A. Whitehead

**Author notes:** E.E.W. Ginkgo Bioworks, L.R.A. McKetta Dept. of Chemical Engineering, University of Texas. Correspondence to: Timothy A. Whitehead, Jenny Smoly Caruthers Biotechnology Building, 3415 Colorado Ave, Room E1B32, Boulder, Colorado 80303.

## Abstract

It is incompletely understood how biophysical properties like protein stability impact molecular evolution and epistasis. Epistasis is defined as specific when a mutation exclusively influences the phenotypic effect of another mutation, often at physically interacting residues. By contrast, nonspecific epistasis results when a mutation is influenced by a large number of non-local mutations. As most mutations are pleiotropic, the *in vivo* folding probability – governed by basal protein stability – is thought to determine activity-enhancing mutational tolerance, which implies that nonspecific epistasis is dominant. However, evidence exists for both specific and nonspecific epistasis as the prevalent factor, with limited comprehensive datasets to validate either claim. Using deep mutational scanning we probe how *in vivo* enzyme folding probability impacts local fitness landscapes. We computationally designed two different variants of the amidase AmiE in which catalytic efficiencies are statistically indistinguishable but the enzyme variants have lower probabilities of folding *in vivo*. Local fitness landscapes show only slight alterations among variants, with essentially the same global distribution of fitness effects. However, specific epistasis was predominant for the subset of mutations exhibiting positive sign epistasis. These mutations mapped to spatially distinct locations on AmiE near the initial mutation or proximal to the active site. Intriguingly, the majority of specific epistatic mutations were codon-dependent, with different synonymous codons resulting in fitness sign reversals. Together, these results offer a nuanced view of how protein folding probability impacts local fitness landscapes, and suggest that transcriptional-translational effects are an equally important determinant as stability in determining evolutionary outcomes.

## Introduction

Understanding the mechanisms of molecular evolution is important to molecular biology, virology, evolutionary biology, and protein engineering. Researchers interested in evolving natural proteins, designing proteins *de novo*, or understanding the extent of contingency on extant proteins must contend with the implicit evolutionary limitations set forth by nature. The challenge, then, is to understand what constrains protein evolution and by what mechanisms. How do these factors interact with one another to alter the frequency of mutations with increased fitness in a given environment, and how do they govern evolvability for new functions?

A particularly important component of evolution is epistasis, or the non-additive combination of mutations (Phillips 2008). Epistasis impacts the rate of evolution and the spectrum of possible evolutionary pathways available to a protein (Bershtein et al. 2006). Epistasis is said to be specific if a mutation exclusively influences the phenotypic effect of other mutations, usually at physically interacting residues (Starr and Thornton 2016). By contrast, nonspecific epistasis results when a mutation impacts a global property like stability that can be rescued by large numbers of non-local mutations. Of the two classes, specific epistatic effects exert the greatest influence on the possible evolutionary outcomes (Starr and Thornton 2016). This is because specific epistatic mutations decrease the reversibility of a protein in an evolutionary trajectory. Specific epistasis also decreases the robustness of a protein to new mutations, reducing the number of possible evolutionary paths available. By modulating the evolutionary trajectories available, epistatic phenomena exert immense influence over the long-term evolution of proteins (Breen et al. 2012).

What is still incompletely understood is how biophysical parameters like protein stability constrain epistasis. Protein stability as defined here is the cumulative balance of the thermodynamic stability, the folding rate, and the fidelity of the association process for oligomeric proteins; these parameters combine to determine the likelihood that an enzyme will assume its native state (Baker and Agard 1994; Shakhnovich 1997). Analyses of the impacts of stability in evolution at the genomic (Jordan et al. 2015), protein (Tokuriki et al. 2006; Campbell et al. 2016; Kumar er al. 2017), and organismal (Serohijos and Shakhnovich 2014) levels have uncovered a complex and dynamic equilibrium between stabilizing and destabilizing mutations. For enzymes in particular, previous studies have shown that missense mutations often act pleiotropically in which catalytically enhancing mutations are, on average, moderately destabilizing (Tokuriki et al. 2008; Tokuriki et al. 2012; Klesmith et al. 2017). Consequently, high basal stability can buffer catalytically beneficial but destabilizing mutations (Bloom et al. 2006; Yu and Dalby 2018a), allowing fixation. Deleterious destabilizing mutations can be repaired by reversion mutations (Ashenberg et al. 2013), or by specific and non-specific epistatic mutations that rescue stability (Jorden et al. 2015; Yu and Dalby 2018b). These epistatic mutations are a central phenomenon in the stabilizing-destabilizing equilibrium, with significant consequences in long-term evolution (Shah et al. 2015; Dasmeh et al. 2018). It is uncertain whether specific or non-specific epistatic mutations are more likely to rescue a destabilized protein, with evidence existing for both arguments (Ashenberg et al. 2013; Yu and Dalby 2018b; Shah et al. 2015; Dasmeh et al. 2018; Pollock et al. 2012).

Deep mutational scanning experiments provide a wealth of mutational data that can be used to address questions in molecular evolution (Fowler et al. 2014). This technology comprises the use of large mutational libraries with selections coupled to deep sequencing to evaluate relative fitness of thousands of variants in a massively parallel fashion (Klesmith et al. 2015, Kowalsky et al. 2015; Klesmith et al. 2017; Wrenbeck et al. 2017). We previously used deep mutational scanning on the homohexameric aliphatic amidase AmiE from *Pseudomonas aeruginosa* to understand how local fitness landscapes, defined here as the set of all possible single-point amino acid substitutions from wild-type (WT), change with different substrates (Wrenbeck et al. 2017). In this original study, AmiE was chosen as a model as it is stable in its genetic background and has a high probability of folding upon translation. To comprehensively assess how the initial probability of folding in vivo constrains mutational outcomes, we designed two variants of AmiE in which catalytic activity is unperturbed but the proteins have different probabilities of folding *in vivo*. We then used deep mutational scanning to probe the local fitness landscapes around these variants. While we found moderate epistasis, local fitness landscapes are largely insensitive to the initial folding probability of the enzyme variant. In particular, the great majority of beneficial mutations were shared between all three starting points. However, positive sign epistasis was present and was dominated by specific epistasis. Remarkably, we found that the sign of the fitness metric for many mutations depends on the codon preference, suggesting more complicated fitness landscapes than predicted from intrinsic protein biophysics. Together, these results provide a nuanced view of how local fitness landscapes are perturbed under slightly different initial folding probabilities.

## Results

The experimental pipeline used in this study is shown in figure 1A. First, we designed variants of AmiE that possess wild-type catalytic activity but with a reduced probability of folding *in vivo*. Second, we developed selection conditions for the variants such that cell growth is proportional to enzyme activity using a growth selection with acetamide as the sole nitrogen source. Third, near-comprehensive single-site saturation mutant libraries for our variants were prepared (Wrenbeck et al. 2017) and growth selections performed. Fourth, pre- and post-selection populations were deep sequenced to extract mutant frequencies in the selected and reference populations. These frequencies were converted into a relative fitness metric (*ζ*_*i*_) for each mutant *i* defined as

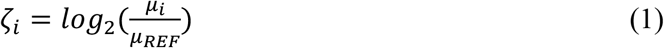

where *μ*_*i*_ and *μ*_*REF*_ represent the specific growth rates in selection media for the mutant (μ_i_) and unmutated AmiE variant (μ_ref_), respectively. A relative fitness score above zero means that a strain harboring a given mutant has higher fitness than those carrying the unmutated variant. It is important to understand the limitations of deep mutational scanning experiments. While these experiments provide quantitative measurements of fitness relative to the respective backgrounds, deep mutational scanning cannot provide information on why a given mutation is beneficial or deleterious. Thus, specifying whether a mutation impacts stability, catalytic efficiency, on-path folding rate, etc. is left to reasoned speculation or further biophysical analysis.

**Fig. 1.**
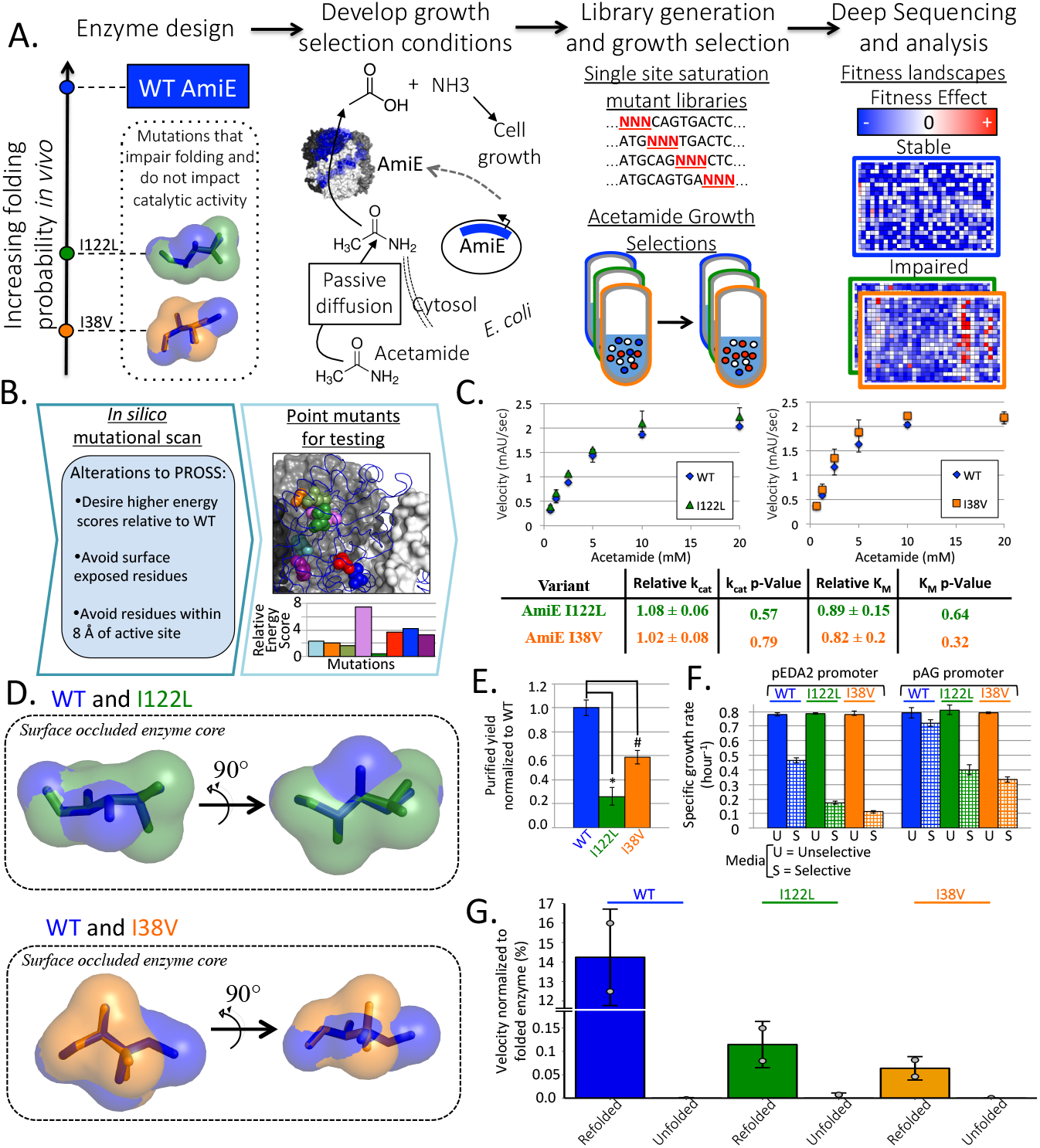
Design of deep mutational scanning experiment. **A.** A graphical overview of this study. Two AmiE enzyme variants with single point mutants (I38V and I122L) with WT catalytic function and lower probabilities of folding *in vivo* were computationally designed and validated experimentally. Constitutive expression of each enzyme from a plasmid was tuned such that the growth rate of our bacterial growth selection strain in selection media was dependent on the expression of functional AmiE. Deep mutational scanning was performed on these variants and compared with WT AmiE. **B-G.** Design and validation of AmiE variants. **B**. A graphical representation of the computational enzyme design. **C**. Enzyme velocity as a function of acetamide and Michaelis-Menten parameters determined relative to WT AmiE. Error bars = 1 s.d., n = 2. **D**. Structural modeling of the cavities introduced into AmiE by designed mutations. **E**. Enzyme yield following *E. coli* auto-induction expression. Error bars = 1 s.d., n = 3, * = p-value = 0.0003, # = p-value = 0.002. **F**. Specific growth rates of strains in M9 (unselective) and in minimal media with 10 mM acetamide as sole nitrogen source (selective) (pEDA2 – low expression, pAG – high expression). Error bars = 1 s.d., n ≥ 3. **G**. Comparison of enzyme reaction velocities at substrate saturation relative to a folded control. Grey dots represent biological replicates, Error bars = 1 s.d., n = 2.

### AmiE variants with lower folding probabilities and wild-type catalytic efficiencies engineered

We first sought to identify mutations to AmiE that, under the selection conditions, would decrease the *in vivo* folding probability of the protein while maintaining wild-type catalytic efficiency. To identify such mutants, we chose to use a computational approach by modifying PROSS (Goldenzweig et al. 2016). Briefly, PROSS designs a protein sequence that will have an improved probability of reaching the folded state *in vivo* relative to its input. This improved folding probability correlates with biophysical properties like improved protein stability, faster on-target folding rate, or reduced aggregation propensity. As our experimental objective is essentially the inverse problem, we modified the Rosetta FilterScan protocol undergirding PROSS and then selected point-mutations with higher energy scores relative to AmiE wild-type (WT) (fig. 1B, **supplementary table S1, Supplementary Material** online). For each mutant these scores were then cross-referenced with experimental relative fitness scores previously determined for AmiE (Wrenbeck et al. 2017) to ensure that their relative fitness was below zero (**supplementary table S1, Supplementary Material** online).

Of thirteen variants with 1-3 mutations from WT selected for experimental characterization, nine expressed as soluble proteins in *E. coli* BL21* (DE3). We purified a subset of these nine variants and assessed their catalytic efficiency with the substrate acetamide. While most mutants showed reduced enzymatic activity, both AmiE I38V and AmiE I122L showed statistically indistinguishable maximum turnover rates (k_cat_) and Michaelis constants (K_M_) compared with WT (fig. 1C, **supplementary table S2, Supplementary Material** online). Furthermore, size exclusion chromatography showed no oligomeric differences between WT and AmiE I38V or AmiE I122L (**supplementary fig. S1, Supplementary Material** online) in PBS at 30 μM, suggesting that the variants maintain the expected homohexameric quaternary structure.

AmiE I38V removes a methyl group to open a small cavity in the core, while I122L modulates hydrophobic core packing in the monomer subunit (fig. 1D, **supplementary fig. S2, Supplementary Material** online). Both mutations are located in the hydrophobic core distal from the dimeric and homohexameric contacts necessary for quaternary assembly (**supplementary fig. S2, Supplementary Material** online). Based on Rosetta analysis, we predict that these mutations disrupt the core of AmiE resulting in thermodynamic destabilization of the native monomer. However, mutations in the stability cores of proteins can disrupt the hierarchy of folding (Raschke and Marqusee 1997) through the destabilization of folding intermediates (Raschke et al. 1999), by limiting the intermediate states accessible during folding (Karshikoff et al. 2015), and by decreasing the thermodynamic stability (Robic et al. 2002) of the monomeric subunits outside of the quaternary structure (Scholl et al. 2015).

To distinguish among these possibilities, we attempted tryptophan fluorescence unfolding measurements using guanidinium-HCl (Gdn-HCl) as a denaturant to determine the effective thermodynamic stability. However, AmiE WT aggregated in moderate Gdn-HCl concentrations under most conditions (data not shown), and under conditions of no aggregation and complete unfolding no isosbestic point was recovered (**supplementary fig. S3, Supplementary Material** online). This lack of an isosbestic point indicates more complicated reversible folding at 4°C than simple 2-state models. We also performed thermal shift assays with the purified homohexameric enzymes in a series of dilutions (10, 5, 1, 0.5, 0.25 μM) (**supplementary fig. S4, Supplementary Material** online) to measure thermal stabilities. Two-state irreversible unfolding curves were obtained at 10 and 5 μM. Analysis of the melting curves reveals statistically indistinguishable melting temperatures between WT and variants (**supplementary table S3, Supplementary Material** online). This data suggests that all have a single transition from homohexamer into unfolded monomers. It is well established that oligomeric proteins are often more stable than in their natively folded monomeric or dimeric forms (Scholl et al. 2015). Thus it is likely that the thermal melt is measuring the stability of homohexameric assembly, which would be expected to be identical between WT and variants as neither mutation resides at an oligomeric interface. To identify AmiE concentrations for which the monomeric form is favored, we reasoned that hexameric dissociation would result in inactive enzyme (Cervoni et al. 2003), which could be measured colorimetrically using our established activity assay. Indeed, activity analysis of dilute solutions of AmiE WT at 500 nM showed larger decreases in activity than 900 nM over moderate incubation periods (**supplementary fig. S5, Supplementary Material** online). Unfortunately, usable signal was not detected at protein concentrations of less than 5 μM in thermal melt assays (**supplementary fig. S4, Supplementary Material** online).

While we were unable to determine thermal or thermodynamic stabilities of the AmiE monomers, complementary *in vitro* and *in vivo* experiments support that both I122L and I38V variants have lower *in vivo* folding probabilities than WT in the general order: I38V<I122L<WT. For both variants, all synonymous mutations had a fitness metric below zero (**supplementary table S1, Supplementary Material** online), suggesting loss of fitness as a result of change at the protein level (Wrenbeck et al. 2017). When driven from the same T7 promoter under identical Studier auto-induction (Studier 2005) protein expression conditions, both AmiE I38V and I122L have statistically significant lower purification yields of soluble protein than WT (fig. 1E and **supplementary table S2, Supplementary Material** online). Furthermore, *E. coli* harboring the AmiE variants expressed from the same plasmid – pEDA2 (Wrenbeck et al. 2017) maintaining the same constitutive promoter, ribosome-binding site (RBS), and 5’ untranslated region (5’ UTR) – showed lower specific growth rates than WT when grown with acetamide as the sole nitrogen source (fig. 1F and **supplementary table S2, Supplementary Material** online). These results suggest that the folding probability upon translation for these variants is lower than for WT. Finally, while denatured WT can refold into active enzyme at 14.2% yield, both I38V (0.06%) and I122L (0.11%) have vastly lower refolding yields (fig. 1G and **supplementary table S2, Supplementary Material** online). These results together support a model where the I38V and I122L mutations result in a lower probability of correctly folding into active homohexameric enzyme *in vivo*.

### Deep mutational scans for AmiE variants

Deep mutational scanning of these variants was performed using a previously developed growth selection (Wrenbeck et al. 2017) in media with 10 mM acetamide as the sole nitrogen source. These growth selections required tuning the constitutive amidase expression such that the specific growth rate of variant *i* expressed in *E. coli* MG1655 *rph+* in the selection media relative to that in defined minimal media (μ_s,i_/μ_M9,i_) is 0.4-0.6. However, plasmid pEDA2 used for AmiE WT selections did not support high enough growth rates for the I38V and I122L variants (fig. 1F and **supplementary table S2, Supplementary Material** online). Thus, we screened additional promoters for AmiE I38V and AmiE I122L while maintaining the same 5’ UTR and RBS for all constructs in order to minimize potential variant-dependent mRNA effects on fitness (**supplementary table S4, Supplementary Material** online). Plasmid pAG with a stronger constitutive promoter than pEDA2 supported a growth rate ratio of 0.49 ± 0.03 for AmiE I122L and 0.42±0.02 for AmiE I38V (fig. 1F and **supplementary table S2, Supplementary Material** online). By contrast, pAG AmiE WT had a nearly 2-fold higher growth rate ratio of 0.91 ± 0.04 (fig. 1F and **supplementary table S2, Supplementary Material** online).

Next, we generated near comprehensive single-site saturation mutant libraries using nicking mutagenesis (Wrenbeck et al. 2016) (full library statistics are shown in **supplementary table S5, Supplementary Material** online). For AmiE I38V mutations at residues 32-44 flanking the site of the disrupting mutation were not constructed, while for AmiE I122L mutations at residues 115-130 and 132 were not made. Plasmids expressing mutant enzyme libraries were electroporated into *E. coli* MG1655 *rph+* under conditions minimizing double transformants. Then, strains harboring AmiE libraries underwent growth selections in replicate with initial population sizes of >6×10^6^ cells for approximately 8 generations at 37°C. A biological replicate for AmiE WT covering residues 171-255 was also performed to compare with previous published results (Wrenbeck et al. 2017). The pre- and post-selection populations were barcoded and deep sequenced. The resulting data was processed using PACT (Klesmith and Hackel 2018) to obtain the relevant fitness metrics for each mutant in the library. The depth of sequencing ranged from 155 to 300-fold coverage for the libraries (**supplementary fig. S6, Supplementary Material** online). In total, we were able to recover the relative fitness for 93.7% and 91.8% of all possible non-synonymous mutants for AmiE I122L and AmiE I38V, respectively (**supplementary table S6, Supplementary Material** online).

To estimate reproducibility, we compared the AmiE WT replicate selections performed here with data from an identical selection experiment performed in Wrenbeck et al. (2017). Correlation coefficients between selections are ≥0.90 (**supplementary fig. S7, Supplementary Material** online), which is comparable to correlation between replicates performed for this work (AmiE I122L – 0.921; AmiE I38V – 0.952) (fig. 2A). Additionally, there was essentially no correlation between relative fitness and pre-selection frequency of a given mutant in the library, (WT AmiE – R = 0.011, AmiE I122L – R = 0.026, AmiE I38V – R = −0.0376) (**supplementary fig. S8, Supplementary Material** online) indicating that pre-selection read counts do not bias the fitness metrics obtained.

**Fig. 2.**
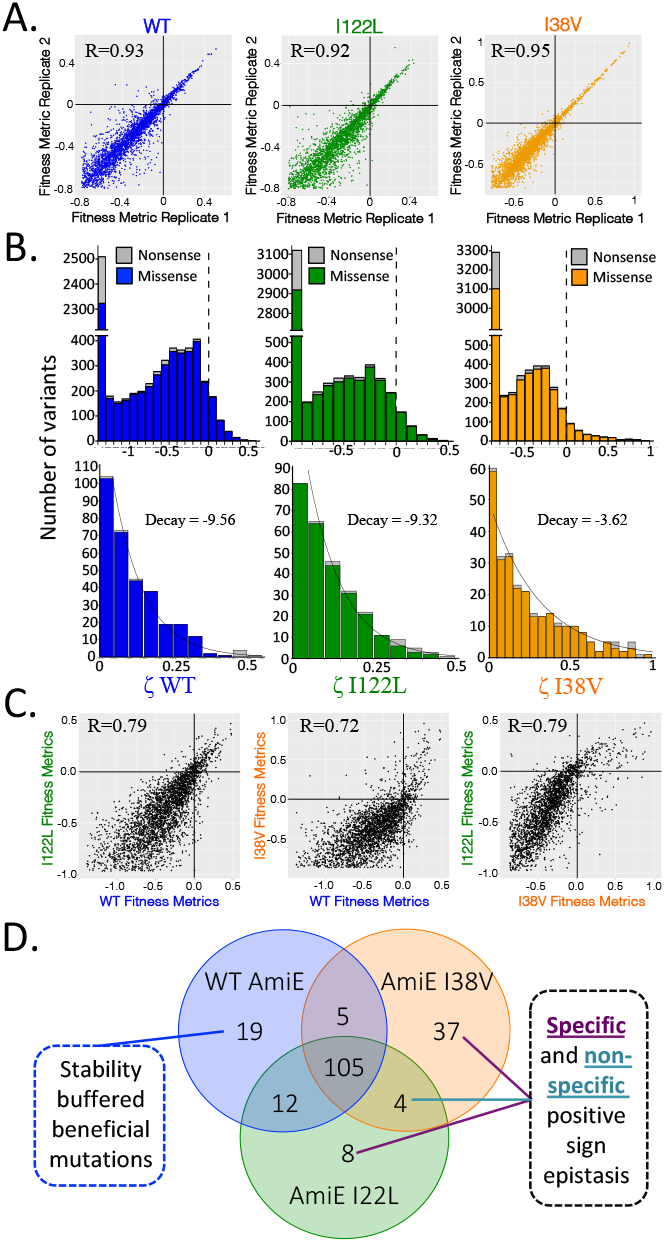
Local fitness landscapes are nearly insensitive to initial protein folding probability *in vivo*. **A.** Correlation between AmiE variant technical replicates. **B**. Distributions of fitness effects (DFE) of nonsense and missense mutations for the AmiE variants. Upper plots show full DFE, while lower plots include only beneficial mutants with best-fit exponential curve. **C**. Correlation of fitness between AmiE variants. **D**. Venn diagram for all unique and shared beneficial mutations for the respective variants.

### Distribution of fitness effects are largely insensitive to initial protein folding probability

The shape of the distribution of fitness effects (DFE) governs the local protein fitness landscape. Realizing that beneficial mutations are rare, the likelihood of finding beneficial mutations was predicted by Orr (2010) to follow the Pareto family of distributions. Using the set of beneficial mutations – variants with relative fitness above wild-type under selective media – we were previously able to describe the shape of the DFE for beneficial mutations as exponential with high statistical power (Wrenbeck et al. 2017). The new datasets allow us to ask directly whether the shape of DFE changes with respect to enzyme folding probability. Consistent with expectations, all variants have very similar distributions of fitness effects (fig. 2B) with a tight range of total possible mutations that are beneficial. For both variants the Pareto family of functions also describes their distributions of beneficial fitness effects (**supplementary table S7, Supplementary Material** online). Thus, given approximately the same relative fitness, the probability of finding rare beneficial mutations is independent of initial likelihood of native folding.

### Moderate epistasis observed with decreasing enzyme folding probability

How does the local fitness landscape change in the background of the AmiE variants? If mutations were completely additive with the disrupting mutations, we would expect the comparison of the local fitness landscapes for the enzymes to have 1:1 correlations and approach the correlation coefficients found between replicates (R ~0.92). On the other hand, complete non-additivity of mutations would lead to minimal correlation. We were able to compare 2,813 mutations above the lower bound of relative fitness (45.4% of possible mutations) shared between the three datasets. Pearson’s correlation analysis of the DFE finds that the WT local fitness landscape is reasonably correlated with that of the variants (WT vs. I38V R= 0.72, WT vs. I122L R = 0.79), and this correlation is similar to that between I122L vs. I38V (R= 0.79) (fig. 2C). Notably, these correlation coefficients are lower than for replicates. Furthermore, linear regression best fits show lower slopes between variants than within replicates (fig. 2A,C). Combined, these results indicate that while in bulk the majority of mutations behave identically in different genetic backgrounds, there may be some epistasis.

Next, we tested for increased negative epistasis. We predicted that the greater folding probability of AmiE WT provides a buffering effect, which could temper many deleterious mutations. To test this hypothesis, we generated empirical cumulative distribution functions (ECDF) for the deleterious mutations for each of the three enzymes (**supplementary fig. S9 and table S8, Supplementary Material** online). This analysis is limited to the range of deleterious mutations quantitatively captured in our experimental system (Kowalsky et al. 2015). Within this range, the application of the Kolmogorov-Smirnoff test led to the acceptance of the null hypothesis that the ECDF of AmiE WT is below that of AmiE I38V (**supplementary table S8, Supplementary Material** online). By the same test, we were unable to discriminate the ECDFs of AmiE WT and AmiE I122L (**supplementary table S8, Supplementary Material** online). Therefore, AmiE I38V has a greater number of more deleterious mutations than AmiE WT. This indicates that the lower folding probability of the AmiE I38V background increases the likelihood of negative epistatic effects when compared to the more resilient AmiE WT.

Precise measurements of negative and positive epistasis are complicated by the relatively narrow range of fitness our experimental system captures. However, we can determine the sign of fitness in our datasets with high precision. To evaluate the relative prevalence of sign epistasis, we defined any mutant as beneficial if *ζ*_*i*_ > 0 for both replicates and if *ζ*_*i*_ > 0 within a 95% confidence interval (see **Materials and Methods**). Conversely, we define a mutant as deleterious if *ζ*_*i*_ < 0 for both replicates and if *ζ*_*i*_ < 0 within a 95% confidence interval. We used these cutoffs to sort beneficial variants into the seven possible fitness bins (fig. 2D). Using a stricter requirement – that beneficial mutants are defined as those with a ≥10% increase in specific growth rate over the genetic background – leads to similar results (**supplementary fig. S10, Supplementary Material** online).

### Most beneficial mutations in WT background are shared

Previous studies found that stable proteins can buffer destabilizing mutations that are otherwise beneficial (Tokuriki and Tawfik 2009a; Gong et al. 2013). We are able to assess the extent of this phenomenon in our datasets. We find that 122/141 (86.5%) of beneficial mutations in the WT background are also beneficial in the I38V and/or the I122L genetic background (fig. 2D). Of these, six globally beneficial mutations (S9A, A28R, R89E, I165C, V201M, A234M) have been previously characterized biophysically (Wrenbeck et al. 2017) and are known to improve specific amidase flux under the selection conditions of 10 mM acetamide at 37°C. Conversely, only 19 of 141 beneficial mutations (13.5%) are specific in the WT background (fig. 2D). Therefore, beneficial mutations that are buffered in the stable background are present but in the minority.

The 19 WT-specific beneficial mutations map to two predominant locations: eleven (G291C, L297H, D311C, E320A, S325Y/T/C, 326Y/H, R336L, G341Y; fig. 3A) are at the extreme C-terminus that creates extensive homodimer contacts, while four (M72T, E74G, A78H, E82I; fig. 3A) are located on helix B and helix C distal from any oligomeric contacts in the homohexamer. The majority of the C-terminal mutations are predicted to destabilize the homodimerization interface, which we speculate could lead to subtle structural rearrangements in the AmiE active site. Probable mechanisms behind the helix B/C mutations are more obscure as three of these mutations are at surface exposed positions over 10 Å away from any active site residue. Nevertheless, these findings indicate localized regions where small-scale mutational perturbations lead to increased fitness in a more stable genetic background.

**Fig. 3.**
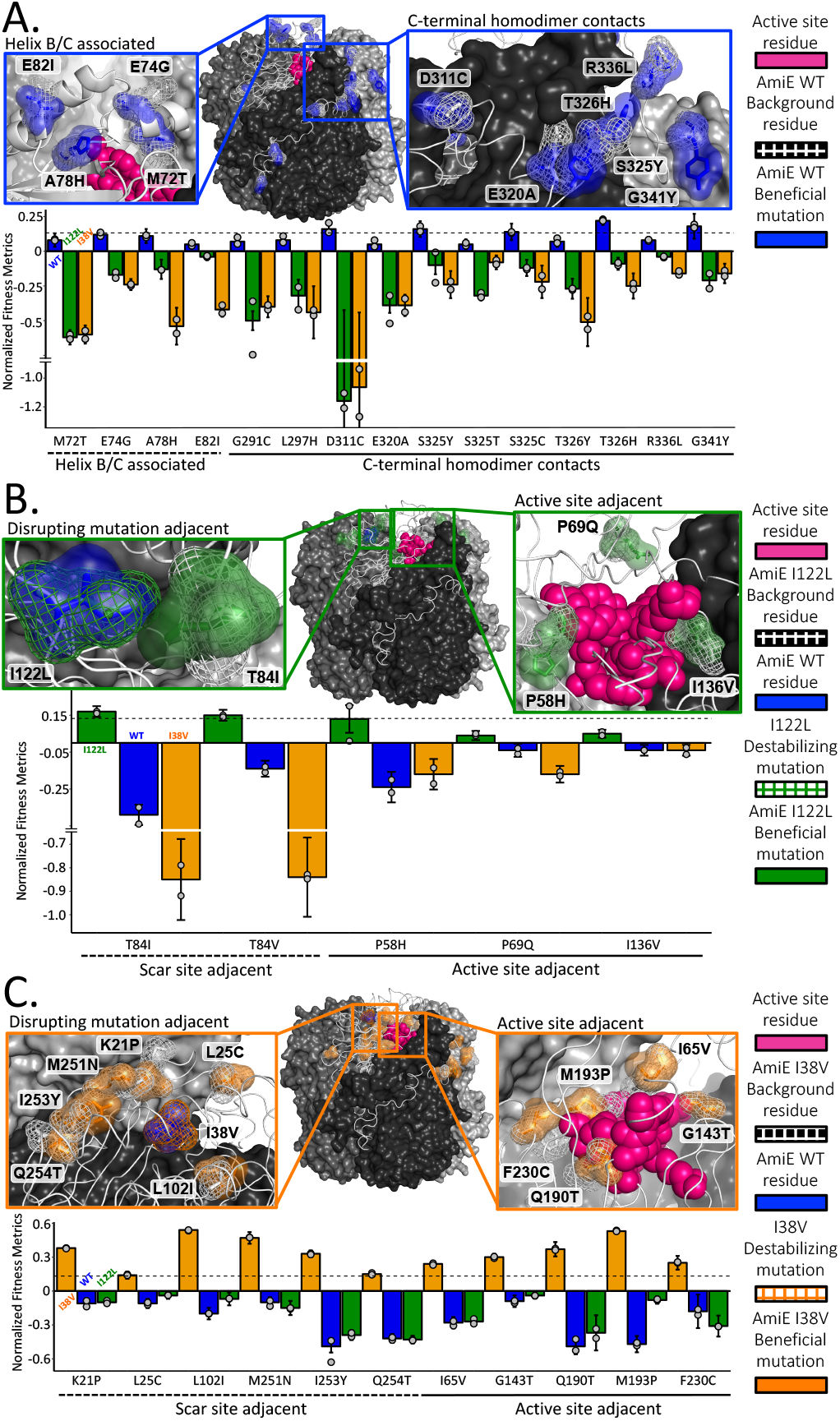
Positive sign epistatic mutations are spatially segregated and specific. **A-C.** Bar graphs of fitness metrics. Error bars represent 95% confidence intervals, while grey dots are fitness metric for each replicate. Horizontal dotted lines represent the cutoff value for mutations that increase the growth rate by ≥10%. Models show: trimer of dimers (wire + surface models), background residues for a respective enzyme (white mesh + sticks), the active site residues (magenta spheres), and where applicable the original mutation (green or orange mesh + sticks). **A**. AmiE WT unique beneficial mutations are located at the C-terminal tail or in the B/C helices. **B.** Location of AmiE I122L unique beneficial mutations segregate to either positions adjacent to position 122 (T84I/V) or adjacent to the active site. **C.** Locations of a subset of AmiE I38V unique beneficial mutations. In **B** and **C** the blue transparent surface with sticks represents the residue in AmiE WT, and green or orange transparent surfaces with sticks represent respective unique beneficial mutations.

To avoid information loss from the simple categorical separation of mutations into bins, normalized linear regression analysis was performed on correlation plots for the beneficial mutations shared in all backgrounds (**supplementary fig. S11, Supplementary Material** online). Correlation for AmiE WT compared with AmiE I38V and AmiE I122L are strikingly similar. In both, the mutational effects are highly disperse and widely deviate from predicted fitness metric function for non-epistatic mutational combinations (Y = 1X + 0). This indicates that non-specific epistatic effects are both present and are imposing large impacts on this population of beneficial mutations.

### Positive sign epistasis is overwhelmingly specific

Our datasets allow direct comparisons between the prevalence of specific and nonspecific epistasis in a folding impaired background. Contrary to previous literature on different model proteins (Bershtein et al. 2006; Bloom et al. 2006; Gong et al. 2013; Huang et al. 1997; Sideracki et al. 2001; Bloom et al. 2005; Tokuriki and Tawfik 2009b), we find that specific reciprocal positive sign epistatic mutations dominate nonspecific mutations in the I38V and I122L backgrounds. In particular, we find 8 specific – unique beneficial – mutations for AmiE I122L and 37 specific mutations for AmiE I38V, compared with only 4 nonspecific – shared beneficial – mutations (fig. 2D and 3B-3C). Analysis of the locations of the unique mutations in the structures of AmiE I122L and AmiE I38V suggest biophysical interpretations of their epistatic mechanism. For AmiE I122L the strongest specific mutations T84I/V directly contact 122L (fig. 3B), while similar mutations (K21P, L25C, L102I, M251N, I253Y, Q254T) occur on loops adjacent to I38V (fig. 3C). Other specific mutations line the enzyme active site. For AmiE I122L (fig. 3B) there are three (P58H, P69Q, I136V), while in AmiE I38V (fig. 3C) there are five (I65V, G143T, Q190T, M193P, F230C). Notably, since our datasets do not include immediately adjacent mutations for I38V and I122L, the extent of specific positive sign epistasis is probably underestimated.

### A plurality of unique beneficial mutations is codon-dependent

Fitness conferred by weak-link enzymes depends on intrinsic protein biophysics but also on mRNA sequence-dependent effects. Perhaps best appreciated of these are synonymous mutations in the first ten codons of a polypeptide because they can substantially alter mRNA stability and access to the ribosome binding sites in bacteria (Wrenbeck et al. 2017, Kristofich et al. 2018). Additionally, synonymous codons can differentially affect cotranslational folding (Buhr et al. 2016) and impact fitness. Here we mapped the variance between synonymous codons encoding beneficial missense mutations (fig. 4A). For all three variants, the majority of high variance codons occur in the first 10 codons, as expected (**supplementary fig. S12A, Supplementary Material** online). However, there were localized punctae of high variance at several downstream positions for both WT and I38V datasets corroborated in replicate measurements. While overall variance in beneficial synonymous codons is weakly or not statistically significant among datasets (**supplementary fig. S12B, Supplementary Material** online), contingency table analysis of synonymous codon fitness disparities for the unique and shared beneficial mutations finds correlation with unique beneficial mutations in the WT and I38V backgrounds (WT p-value = 2.5×10^−8^, I122L p-value = 0.053, I38V p-value = 0.00017; 2-tailed Fisher exact probability test) (fig. 4B). In fact, the vast majority of WT (84.2%) – and many of the I122L (42.8%) and I38V (48.6%) – unique beneficial mutations have synonymous codon fitness sign disparities (fig. 4B). This indicates that transcriptional-translational effects impose significant evolutionary constraints.

**Fig. 4.**
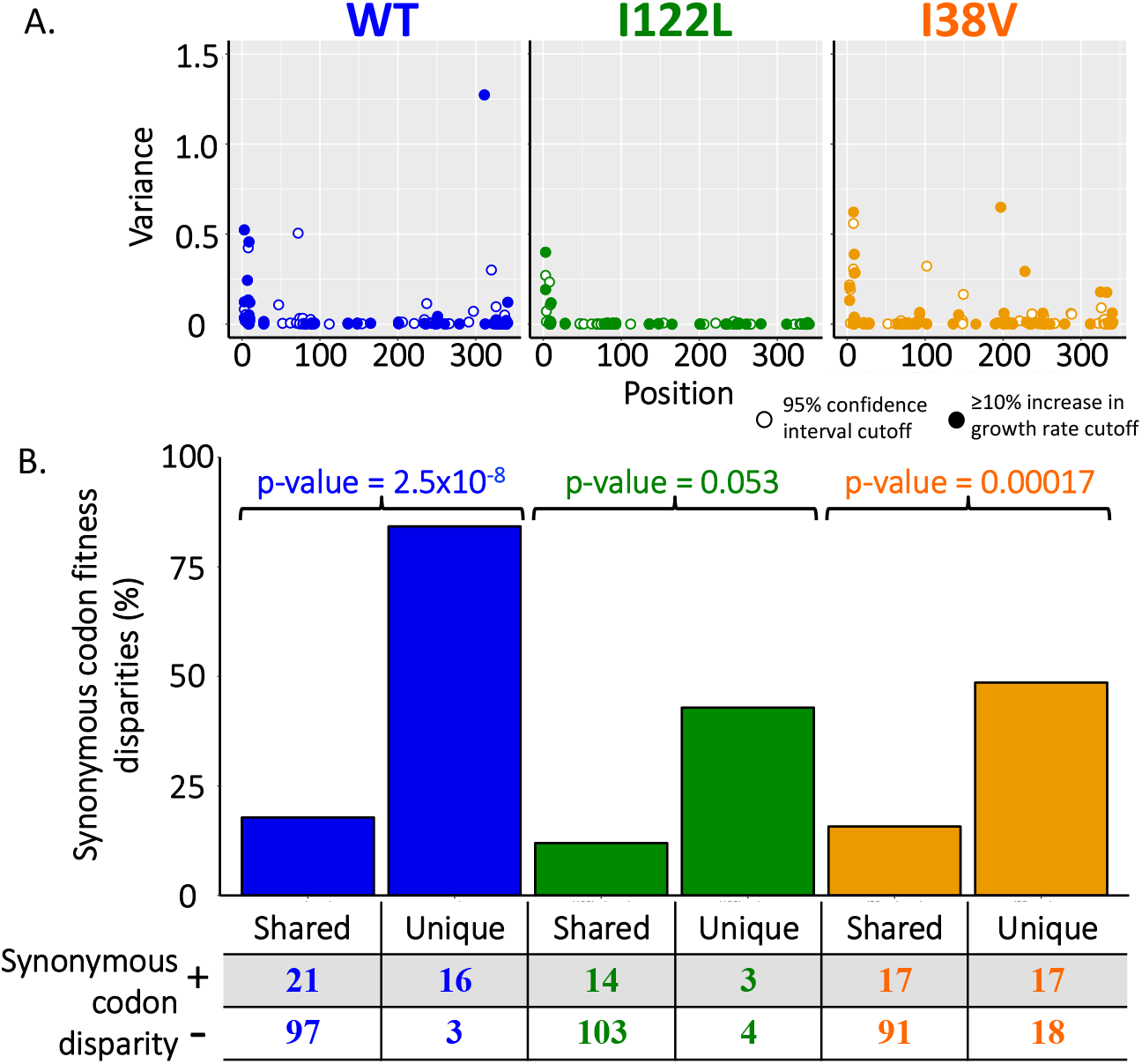
Unique beneficial mutations have high percentages of synonymous codon fitness disparities. **A.** Variance of fitness metrics for synonymous codons of beneficial mutations as a function of position in the primary sequence. **B.** Percentage of shared and unique beneficial mutations with synonymous codon fitness metric disparities. p-values reported are from contingency table analysis with 2-tailed Fisher exact probability test.

## Discussion

In this study we used deep mutational scanning to analyze how the probability of attaining the folded, active state impacts local fitness landscapes. AmiE I38V and AmiE I122L were designed and validated to have identical catalytic parameters to WT but have a lower probability of folding under selection conditions. We found that the DFE for both variants was largely similar to the AmiE WT, and that most fitness-enhancing mutations are shared. However, there were two major surprises found when analyzing the set of mutations exhibiting positive sign epistasis.

First, we expected there to be a larger subset of beneficial mutations shared only between the I122L and I38V datasets, as current models of stability-induced epistasis posit that many nonspecific globally-distributed mutations can improve the probability of folding (Starr and Thornton 2016). In contrast, we found that sign epistatic mutations were overwhelmingly specific for the I122L and I38V backgrounds. It is possible that beneficial non-specific sign epistatic mutations are the minority because of the alterations to *in vivo* expression that are required for the deep mutational scans. In the non-promoter tuned *E. coli* the impaired enzymes provide very weak cell growth. It is possible that non-specific epistatic effects may have arisen to be equivalent with, or dominant to, specific epistatic phenomenon if experiments could be performed without increasing the *in vivo* expression of the impaired enzymes. A limitation of our deep mutational scanning experiments is that cell growth must be proportional to enzyme function and must operate within a window of growth rate ratios (Kowalsky et al. 2015). This requires increased *in vivo* expression of the impaired enzymes. By increasing the *in vivo* expression it is possible that the selective advantages that the non-specific mutations might possess could be dampened. Therefore, the sparsity of non-specific beneficial mutations could be an artifact of our experiments. Further, more careful measurements on a wider array of oligomeric proteins should resolve this seeming contradiction.

As a second surprise, we found that unique beneficial mutations strongly depend on codon choice, as approximately 50% of sign epistatic mutations in the I38V background show sign disparities. We speculate that this unexpected result arises from the complicated co-translational folding *in vivo* of the homohexameric AmiE. Local, specific nonsynonymous mutations may recover on-target folding trajectories more efficiently than nonspecific, globally stabilizing mutations. Similarly, on-pathway folding kinetics may differ considerably between variants, which can be selectively modulated by codon choice. As an alternative explanation, Kudla and colleagues recently report that synonymous codons can exert fitness effects through RNA toxicity itself (Mittal et al. 2018) through an unknown mechanism.

While we were able to determine that both I122L and I38V mutations decrease the probability of active AmiE expression at 37°C, we were unable to measure the relative stability of the monomeric proteins. For the thermodynamic studies, under most conditions AmiE aggregated, and where suitable conditions were uncovered the lack of an isosbestic point hampered analysis. The thermal melts showed a single transition, most likely due to the hexameric dissociation. Lowering the AmiE concentration supported monomer formation at 25^°^C but with the expense of too low of a signal for analysis of monomer thermal stability. Nevertheless, our results show that simple biophysical models currently used to model protein evolution are incomplete and that biophysical models may need to use kinetic models to account for the folding probability *in vivo*.

## Materials and Methods

### Reagents

All antibiotics were purchased from GoldBio and all purchased enzymes were from New England Biolabs. All other chemicals were purchased from Sigma-Aldrich. Primers and mutagenic oligos were purchased from Integrated DNA Technologies and were designed using either Benchling (www.benchling.com) or the Agilent QuikChange Primer Design Program (www.agilent.com).

### Computational design of folding impaired mutants

The FilterScan Rosetta script (Goldenzweig et al. 2016) was modified to predict mutations that would decrease thermodynamic stability without altering catalytic efficiency. The structural coordinates for AmiE (Andrade et al. 2007) (PDB: 2UXY, 341 residues per monomer) were taken from the Protein Data Bank and prepped for use in Rosetta scripts through the ‘clean_pdb_keep_ligand.py’ script released with Rosetta 3 (Leaver-Fay et al. 2011). The crystal structure data was refined through the “refine.xml” Rosetta scripts (unaltered) from Goldenzweig et al. (2016). To avoid impacting catalytic efficiency residues within 8 Å of the active site were excluded from the FilterScan protocol. Additionally, surface residues were predicted and excluded from computational testing to avoid disturbing the native homohexameric state. The FilterScan script was modified to remove the input from the position specific scoring matrix (PSSM) evolutionary conservation term. Mutants with scores that predicted destabilization of the enzyme and were also shown to decrease relative fitness in a previous study (Wrenbeck et al. 2017) were selected for biophysical analysis.

### Plasmid Construction

The variants selected for biophysical analysis were constructed by mutating the AmiE WT sequence using the single mutation protocol of Nicking Mutagenesis (Wrenbeck et al. 2016). The pEDA3 constitutive expression plasmid from Wrenbeck et al. (2016) was used as the vector for the mutagenesis. Variants were subcloned from the pEDA3 background into protein expression plasmid pET-29b(+) (Novagen) or into the constitutive expression plasmid pEDA2 (Wrenbeck et al. 2017) at *NdeI* and *XhoI* sites using classic restriction cloning. Plasmid pAG was constructed by mutating the −10 and −35 promoter regions of the pEDA3 plasmid using the multi-site nicking mutagenesis protocol from Wrenbeck et al. (2016). Following mutagenesis, the mutant promoter libraries were transformed into the *E. coli* growth selection strain MG1655 rph+ [F-λ-] (Coli Genetic Stock Center #7925, CGSC strain designation: BW30270). All AmiE variant DNA and protein sequences are listed in Supplementary Notes S1 and S2.

### Near Comprehensive single-site mutant library construction

Near comprehensive mutant libraries for AmiE I38V and AmiE I122L were constructed using the comprehensive nicking mutagenesis protocol from Wrenbeck et al. (2016). The genes for both impaired enzymes were broken into the following tiles: tile 1 (residues 1-85), tile 2 (residues 86-170), tile 3 (residues 171-255), and tile 4 (residues 256-341) in pAG. For AmiE I38V 13 residues were excluded from mutagenesis (residues 32-44), while for AmiE I122L 17 residues were excluded from mutagenesis (residues 115-130 and 132). Nicking mutagenesis products were amplified and purified as in Klesmith et al. (2015). 10 ng each of the respective DNA libraries was transformed into *E. coli* MG1655 *rph*+ by electroporation performed with either a 1 mm electroporation cuvette at 1200 V (AmiE I122L; AmiE WT), or a 2 mm electroporation cuvette at 1600 V (AmiE I38V) using an Eppendorf Eporator. All experimental and control libraries were transformed into the selection strain with greater numbers than that required for theoretical complete library coverage (**supplementary table S5, Supplementary Material** online). The transformation procedure for the selection strain was optimized to minimize double plasmid transformants as described in Kowalsky et al. (2015). −80°C freezer cell stocks of the libraries were prepared as detailed in Klesmith et al. (2015).

### Protein expression and purification

pET29(b) constructs harboring genes encoding AmiE variants were transformed into *E. coli* BL21*(DE3) cells (Invitrogen) and expressed using Studier auto-induction (Studier 2005) at 22°C for 16-18 hours. Cultures were pelleted and frozen at −80°C. Proteins were purified from cell pellets by Ni-NTA affinity chromatography exactly as described in Klesmith et al. (2015). Purified enzymes were desalted into phosphate buffered saline (PBS; 10 mM Na_2_HPO_4_, 1.8 mM KH_2_PO_4_, 2.7 mM KCl, 137 mM NaCl, pH 7.4) using disposable PD-10 desalting columns (GE Healthcare), sterilized through a 0.22 µm syringe filter, and stored at 4°C until analysis. Purified enzymes showed as a single band by SDS-PAGE (**supplementary fig. S13, Supplementary Material** online). Purified enzyme solutions were quantified using the absorbance at 280 nm in 1x PBS using a published theoretical A_280_ molar extinction coefficient of 56,980 M^−1^ cm^−1^ (Bienick et al. 2014).

For quantitative comparison of purified product yields under the T7 promoter system in BL21* *E. coli*, slight alterations to the above induction scheme were used. For this analysis induction cultures were always started at an OD_600_ of 0.005 from 1 mL LB + 50 µg/mL kanamycin cultures grown at 37°C overnight. Next, 500 mL induction cultures were inoculated at an OD_600_ of 0.005 with the overnight cultures and grown at 37°C with shaking at 250x rpm for 6 hours, and following this growth step cultures were moved to 22°C and induced for ~17 hours. Induction cultures were then pelleted and the wet cell weights of the pellets recorded. Cultures were then purified, desalted, and quantified as described above.

### Biophysical analysis of proteins

Analysis of the growth rates of the AmiE variants in the respective constitutive expression plasmids in MG1655 rph+ *E. coli* were performed exactly as in Wrenbeck et al. (Wrenbeck et al. 2017). Assessment of the oligomeric state of the purified enzymes was performed using SEC-FPLC. Approximately 3 mL of 30 µM of the purified enzymes in PBS were run on an AKTA-FPLC system at 1 mL/min on an HiLoad 16/600 Superdex 200 column equilibrated with PBS. Enzyme kinetics (K_M_ and k_cat_) was determined via phenol-alkaline hypochlorite end-point activity assays exactly as in Wrenbeck et al. (Wrenbeck et al. 2017). Kinetic analysis of purified WT and folding impaired AmiE variants was performed within 6 days of purification.

To test for an isosbestic point the AmiE variants were denatured in guanidinium-HCl (GDN-HCl) and native tryptophan fluorescence was detected. Purified AmiE WT was diluted to 52 μM in increasing amounts of ice cold GDN-HCl (0 – 4 M) in PBS with 1 mM DTT and mixed gently. 200 μL of the denaturation mixtures were placed into opaque black 96 well plates and covered with optical film. Plates containing the samples were incubated at 4°C for 8 hours. Next, the native tryptophan fluorescence was measured in uncovered plates using an excitation of 290nm and emission of was detected over a range of wavelengths: 310 nm – 370 nm. This lack of an isosbestic point indicates an unfolding model that is more complex than a two-state model, presumably because of monomer folding and homohexamer association.

Thermal melt analysis was performed exactly as described in Wrenbeck et al. (2017) with incubations for 2.5 hours at 25°C prior to addition of the dye and initiation of the thermal shift assays; the data was processed as in Huynh et al. (2015). In brief 10, 5, 2.5, 1, 0.5, and 0.25 μM purified AmiE in 1x PBS was incubated at 25°C for 2.5 hours. Next, the samples had 5 μL of 200x SYPRO-orange dye (Life Technologies) added to 45 μL of the diluted enzymes in 0.1 mL MicroAmp^®^ 96-Well Reaction Plate (Life Technologies) and covered with MicroAmp^™^ optical film (Life Technologies). Thermal melt analysis was performed in a QuantStudio 6 Flex RT-PCR device (ThermoFisher). The melt ranged from 25°C to 98°C with 1°C change per minute and with a 2 minute incubation at the first and last temperatures. Boltzmann curve fitting was performed to determine T_m_ using GraphPad Prism (www.graphpad.com).

To assess relative activity of the refolded enzymes, enzymes were first denatured in ice cold 3 M GDN-HCl in PBS supplemented with 1 mM DTT for 16 hours at 4°C at a final concentration of 50 μM. The solution was then diluted 50-fold into PBS with 1 mM DTT and 0.1% (w/v) BSA at 4°C in 96-well PCR plates that had been blocked with 1% (w/v) BSA in PBS for 1 hour at 37°C, resulting in a total protein concentration of 1 μM. Refolding mixtures were then incubated at 4°C for 5 minutes, and warmed to 37°C over 4.5 minutes. Samples were then held at 37°C for 20 minutes and then cooled to 4°C and held there until assaying. Immediately prior to assaying, samples were diluted in fresh ice-cold PBS with 1 mM DTT to ensure linearity in the activity assays. Enzymes were assayed using the phenol-alkaline hypochlorite end-point assay with 20 mM acetamide as the substrate. As a control, enzymes went through the same steps as above except without initial GDN-HCl denaturation. The percent activity of the refolded enzyme was determined relative to the control sample. Two biological replicates were performed for all reactions. Refolding experiments were performed within 7 days of purification of the enzymes from the pellets.

### Growth selections

Growth selections were performed exactly as in Wrenbeck et al. (Wrenbeck et al. 2017). Briefly, starter cultures were grown overnight in the non-selective media (M9 minimal media: 47.6 mM Na_2_HPO_4_, 22 mM KH_2_PO_4_, 8.54 mM NaCl, 18.68 mM NH_4_Cl, 50 µg/mL carbenicillin, pH 7.0) and the following day the cells were washed in ice-cold M9 salt solution without ammonium chloride. Next, 3 mL of non-selective or selective media (M9 minimal media without ammonium chloride supplemented with 10 mM acetamide) was inoculated with the washed cells at an initial OD_600_ = 0.02 (~6×10^6^ cells) in Hungate tubes. Cultures were grown at 37°C with shaking at 250x rpm for approximately 8 generations. Continuous exponential growth was ensured by harvesting cells after the first 4 generations and re-inoculating with 3 mL fresh media + antibiotic at OD_600_ = 0.02 prior to growth for the final 4 generations. Following 8 generations of growth the cells were stored and plasmid DNA extracted as in Klesmith et al. (2016). Unique selections started from the same unselected overnight culture were performed as replicates. To evaluate reproducibility between growth selections performed here and previous work performed on AmiE WT (Wrenbeck et al. 2017), a deep mutational scan of residues 171-255 in AmiE WT was performed in parallel to each growth selection performed in the present work.

### Deep mutational scanning

AmiE variant DNA collected from the pre- and post-selection libraries was prepared for 300 BP paired end Illumina MiSeq sequencing as in Kowalsky et al. (2015). Primers used to the PCR reactions in preparation for Illumina sequencing are listed in (**supplementary table S9, Supplementary Material** online). Sequencing of the variants was performed at the University of Illinois Chicago sequencing core. AmiE WT deep mutational scanning unprocessed sequencing results from Wrenbeck et al. (Wrenbeck et al. 2017) were downloaded from the SRA. All data was processed using PACT (Klesmith et al. 2018) with the following changes from the default options entered into the configuration file: fast_filter_translate: qaverage = 20, and qlimit = 0; enrichment: ref_count_threshold = 5, sel_count_threshold = 0, strict_count_threshold = True. Normalized fitness metrics (ζ_i_) were calculated by PACT as outlined in Kowalsky et al. (2015). To summarize, PACT calculates an enrichment ratio (ε_i_) for mutations by assessing the pre- and post-selection counts of each mutant:

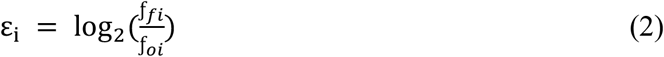

Where *f*_*fi*_ is the frequency of mutant *i* in the post-selection population and *f*_oi_ is the frequency in the pre-selection population. The normalized fitness metric for each mutant i (ζ_i_) was next calculated using the population-averaged number of doublings during selection (g_p_) and the enrichment ratios of the mutant (ε_i_) and the unmutated starting variant (ε_ref_):

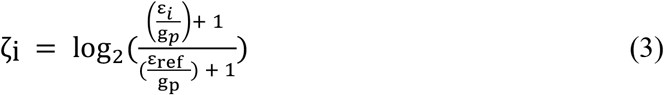

Lower bound fitness metrics – the cut-off below which fitness metrics cannot be discriminated from one another – were calculated as in Wrenbeck et al. (2017) by using the median read count for the pre-selection library and other statistics produced by PACT (**supplementary fig. S6, Supplementary Material** online). Lower fitness metrics were calculated to be: −1.38 for AmiE WT, −0.97 for AmiE I122L, and −0.86 for AmiE I38V.

AmiE I122L had seven outlier mutations removed because the difference in fitness metrics between replicates was greater than the 99.977% confidence intervals determined from sequencing depth of coverage (**supplementary fig. S14, Supplementary Material** online). No other mutants were removed from any of the datasets.

To account for global differences in fitness effects, we also generated normalized scatter plots for globally beneficial mutations. For each pairing of protein variants with AmiE WT (WT/I38V, WT/I122L), we performed a linear regression with the fitness of the first variant as the independent variable and the fitness of the second variant as the dependent variable. We then inverted the linear transformation obtained and applied it to the second fitness. Specifically, if the fitness of the second variant Y was modeled as Y = mX + B, then the normalized fitness Y_hat was computed as (Y-B)/m. Thus, after normalization, the least-squares regression associated with each pairwise plot has a slope of one and an intercept of zero.

### Data Availability

Raw sequencing reads have been deposited in the Sequencing Read Archive (SRA <u>SAMN11258744 – SAMN11258771</u>). Processed data from sequencing reads are given as supplementary information in the Faber_EnzymeStability_ProcessedDatasets excel workbook. All plasmids described in this work have been deposited in AddGene.

## Supporting information

Supporting information part 1

Supporting information part 2

## Acknowledgements

Thanks to Dr. J. Klesmith for his PACT troubleshooting help, E. Maurer and J. Hosten for their help with assorted tasks, and members of the Whitehead lab for providing feedback on ideas and figures. This work was supported by NSF CBET Career Award #1254238 to T.A.W.

