## Supporting information part 1 for "Impact of in vivo protein folding probability on local fitness landscapes"

**Corresponding author: Timothy A. Whitehead**

**This PDF file includes:**

Notes S1 and S2

Figs. S1 to S7

**Other supplementary materials for this manuscript include the following:**

Faber_EnzymeStability_ProcessedDatasets.xls

### Supplementary Notes:

**Note S1: Amino acid sequences of the AmiE variants.** Mutations are highlighted in red.

>AmiE_WT

MRHGDISSSNDTVGVAVVNYKMPRLHTAAEVLDNARKIAEMIVGMKQGLPGM
DLVVFPEYSLQGIMYDPAEMMETAVAIPGEETEIFSRACRKANVWGVFSLTGER
HEEHPRKAPYNTLVLIDNNGEIVQKYRKIIPWCPIEGWYPGGQTYVSEGPKGMKI
SLIICDDGNYPEIWRDCAMKGAELIVRCQGYMYPAKDQQVMMAKAMAWANNC
YVAVANAAGFDGVYSYFGHSAIIGFDGRTLGECGEEEMGIQYAQLSLSQIRDAR
ANDQSQNHLFKILHRGYSGLQASGDGDRGLAECPFEFYRTWVTDAEKARENVE
RLTRSTTGVAQCPVGRLPYEGLEHHHHHH

>AmiE_I122L

MRHGDISSSNDTVGVAVVNYKMPRLHTAAEVLDNARKIAEMIVGMKQGLPGM
DLVVFPEYSLQGIMYDPAEMMETAVAIPGEETEIFSRACRKANVWGVFSLTGER
HEEHPRKAPYNTLVLLDNNGEIVQKYRKIIPWCPIEGWYPGGQTYVSEGPKGMKI
SLIICDDGNYPEIWRDCAMKGAELIVRCQGYMYPAKDQQVMMAKAMAWANNC
YVAVANAAGFDGVYSYFGHSAIIGFDGRTLGECGEEEMGIQYAQLSLSQIRDAR
ANDQSQNHLFKILHRGYSGLQASGDGDRGLAECPFEFYRTWVTDAEKARENVE
RLTRSTTGVAQCPVGRLPYEGLEHHHHHH

>AmiE_I38V

MRHGDISSSNDTVGVAVVNYKMPRLHTAAEVLDNARKVAEMIVGMKQGLPGM
DLVVFPEYSLQGIMYDPAEMMETAVAIPGEETEIFSRACRKANVWGVFSLTGER
HEEHPRKAPYNTLVLIDNNGEIVQKYRKIIPWCPIEGWYPGGQTYVSEGPKGMKI
SLIICDDGNYPEIWRDCAMKGAELIVRCQGYMYPAKDQQVMMAKAMAWANNC
YVAVANAAGFDGVYSYFGHSAIIGFDGRTLGECGEEEMGIQYAQLSLSQIRDAR
ANDQSQNHLFKILHRGYSGLQASGDGDRGLAECPFEFYRTWVTDAEKARENVE
RLTRSTTGVAQCPVGRLPYEGLEHHHHHH

**Note S2: DNA sequences of the AmiE variants.** Codons mutated are underlined and bold, while mutations are highlighted in red.

>AmiE_WT

ATGAGACATGGCGATATTAGCTCGTCAAATGATACCGTAGGCGTAGCCGTGG
TGAATTACAAGATGCCGCGTTTACATACTGCTGCTGAAGTCCTGGATAATGCC
CGCAAAATTGCGGAAATGATCGTTGGTATGAAGCAAGGTCTGCCGGGCATGG
ATCTGGTTGTGTTTCCTGAATATTCTTTACAGGGTATTATGTACGACCCTGCTG
AAATGATGGAAACAGCCGTGGCGATTCCAGGCGAAGAAACGGAAATCTTTA
GCCGTGCTTGTAGAAAAGCAAATGTTTGGGGTGTGTTCTCCCTGACCGGCGA
ACGTCATGAAGAACACCCTAGAAAGGCACCATACAACACTCTGGTCTTGATC
GATAACAACGGTGAAATCGTACAAAAGTACAGAAAGATCATCCCATGGTGTC
CGATTGAAGGCTGGTATCCAGGTGGCCAGACATACGTCTCTGAAGGTCCGAA
AGGCATGAAGATCTCATTAATTATCTGCGATGACGGTAATTATCCGGAAATTT
GGAGAGATTGTGCCATGAAGGGTGCGGAATTGATCGTTCGCTGCCAAGGCTA
TATGTACCCTGCTAAAGACCAACAAGTTATGATGGCTAAGGCAATGGCCTGG
GCGAATAACTGTTATGTCGCTGTAGCAAACGCTGCAGGTTTTGATGGCGTTTA
TAGCTACTTCGGTCATAGTGCCATTATCGGTTTTGACGGCCGTACTCTGGGTG
AATGCGGCGAAGAAGAAATGGGCATTCAATACGCGCAGTTGTCTCTGTCACA
AATCCGCGATGCCCGTGCGAATGACCAAAGTCAGAACCATTTGTTTAAAATC
TTGCACAGAGGTTACTCCGGTTTGCAGGCTTCGGGCGATGGCGACCGTGGTCT
GGCAGAATGCCCATTTGAATTCTACCGTACCTGGGTTACTGATGCTGAAAAG
GCAAGAGAAAACGTGGAACGCCTGACTCGCTCCACAACAGGTGTCGCCCAAT
GCCCAGTAGGTCGTCTGCCGTATGAAGGCCTCGAGCACCACCACCACCACCA
C

>AmiE_I122L

ATGAGACATGGCGATATTAGCTCGTCAAATGATACCGTAGGCGTAGCCGTGG
TGAATTACAAGATGCCGCGTTTACATACTGCTGCTGAAGTCCTGGATAATGCC
CGCAAAATTGCGGAAATGATCGTTGGTATGAAGCAAGGTCTGCCGGGCATGG
ATCTGGTTGTGTTTCCTGAATATTCTTTACAGGGTATTATGTACGACCCTGCTG
AAATGATGGAAACAGCCGTGGCGATTCCAGGCGAAGAAACGGAAATCTTTA
GCCGTGCTTGTAGAAAAGCAAATGTTTGGGGTGTGTTCTCCCTGACCGGCGA
ACGTCATGAAGAACACCCTAGAAAGGCACCATACAACACTCTGGTCTTG**CTC**
GATAACAACGGTGAAATCGTACAAAAGTACAGAAAGATCATCCCATGGTGTC
CGATTGAAGGCTGGTATCCAGGTGGCCAGACATACGTCTCTGAAGGTCCGAA
AGGCATGAAGATCTCATTAATTATCTGCGATGACGGTAATTATCCGGAAATTT
GGAGAGATTGTGCCATGAAGGGTGCGGAATTGATCGTTCGCTGCCAAGGCTA
TATGTACCCTGCTAAAGACCAACAAGTTATGATGGCTAAGGCAATGGCCTGG
GCGAATAACTGTTATGTCGCTGTAGCAAACGCTGCAGGTTTTGATGGCGTTTA
TAGCTACTTCGGTCATAGTGCCATTATCGGTTTTGACGGCCGTACTCTGGGTG
AATGCGGCGAAGAAGAAATGGGCATTCAATACGCGCAGTTGTCTCTGTCACA

AATCCGCGATGCCCGTGCGAATGACCAAAGTCAGAACCATTTGTTTAAAATC
TTGCACAGAGGTTACTCCGGTTTGCAGGCTTCGGGCGATGGCGACCGTGGTCT
GGCAGAATGCCCATTTGAATTCTACCGTACCTGGGTTACTGATGCTGAAAAG
GCAAGAGAAAACGTGGAACGCCTGACTCGCTCCACAACAGGTGTCGCCCAAT
GCCCAGTAGGTCGTCTGCCGTATGAAGGCCTCGAGCACCACCACCACCACCA
C

>AmiE_I38V

ATGAGACATGGCGATATTAGCTCGTCAAATGATACCGTAGGCGTAGCCGTGG
TGAATTACAAGATGCCGCGTTTACATACTGCTGCTGAAGTCCTGGATAATGCC
CGCAAA**GTT**GCGGAAATGATCGTTGGTATGAAGCAAGGTCTGCCGGGCATGG
ATCTGGTTGTGTTTCCTGAATATTCTTTACAGGGTATTATGTACGACCCTGCTG
AAATGATGGAAACAGCCGTGGCGATTCCAGGCGAAGAAACGGAAATCTTTA
GCCGTGCTTGTAGAAAAGCAAATGTTTGGGGTGTGTTCTCCCTGACCGGCGA
ACGTCATGAAGAACACCCTAGAAAGGCACCATACAACACTCTGGTCTTGATC
GATAACAACGGTGAAATCGTACAAAAGTACAGAAAGATCATCCCATGGTGTC
CGATTGAAGGCTGGTATCCAGGTGGCCAGACATACGTCTCTGAAGGTCCGAA
AGGCATGAAGATCTCATTAATTATCTGCGATGACGGTAATTATCCGGAAATTT
GGAGAGATTGTGCCATGAAGGGTGCGGAATTGATCGTTCGCTGCCAAGGCTA
TATGTACCCTGCTAAAGACCAACAAGTTATGATGGCTAAGGCAATGGCCTGG
GCGAATAACTGTTATGTCGCTGTAGCAAACGCTGCAGGTTTTGATGGCGTTTA
TAGCTACTTCGGTCATAGTGCCATTATCGGTTTTGACGGCCGTACTCTGGGTG
AATGCGGCGAAGAAGAAATGGGCATTCAATACGCGCAGTTGTCTCTGTCACA
AATCCGCGATGCCCGTGCGAATGACCAAAGTCAGAACCATTTGTTTAAAATC
TTGCACAGAGGTTACTCCGGTTTGCAGGCTTCGGGCGATGGCGACCGTGGTCT
GGCAGAATGCCCATTTGAATTCTACCGTACCTGGGTTACTGATGCTGAAAAG
GCAAGAGAAAACGTGGAACGCCTGACTCGCTCCACAACAGGTGTCGCCCAAT
GCCCAGTAGGTCGTCTGCCGTATGAAGGCCTCGAGCACCACCACCACCACCA
C

### Supplementary Figures:

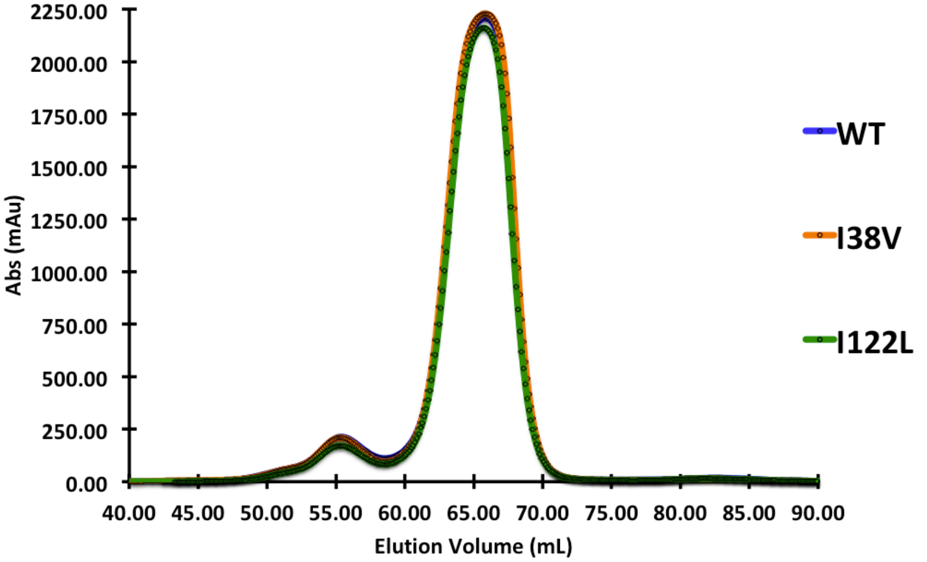


**Fig. S1: Chromatogram of purified AmiE variants.** SEC-FPLC chromatograms of AmiE proteins run on a HiLoad 16/600 Superdex 200 column at 1 mL/min in PBS as the mobile phase. No gross differences in oligomeric state were determined for the proteins.

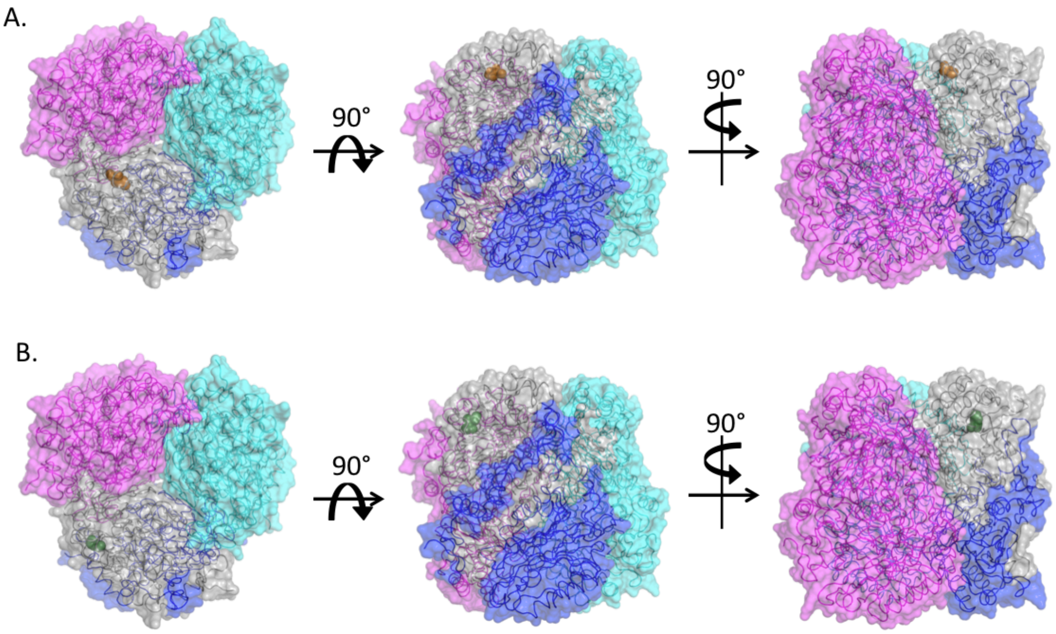


**Fig. S2: Locations of I38V and I122L mutations in AmiE quaternary structure**. A. Three perspectives on the location of the disrupting I38V mutation, the residue I38 is shown as orange spheres. B. Three perspectives on the location of the disrupting I122L mutation, the residue I122 is shown as green spheres. Both mutations are in the monomer core and are not be predicted to impact quaternary structure or assembly.

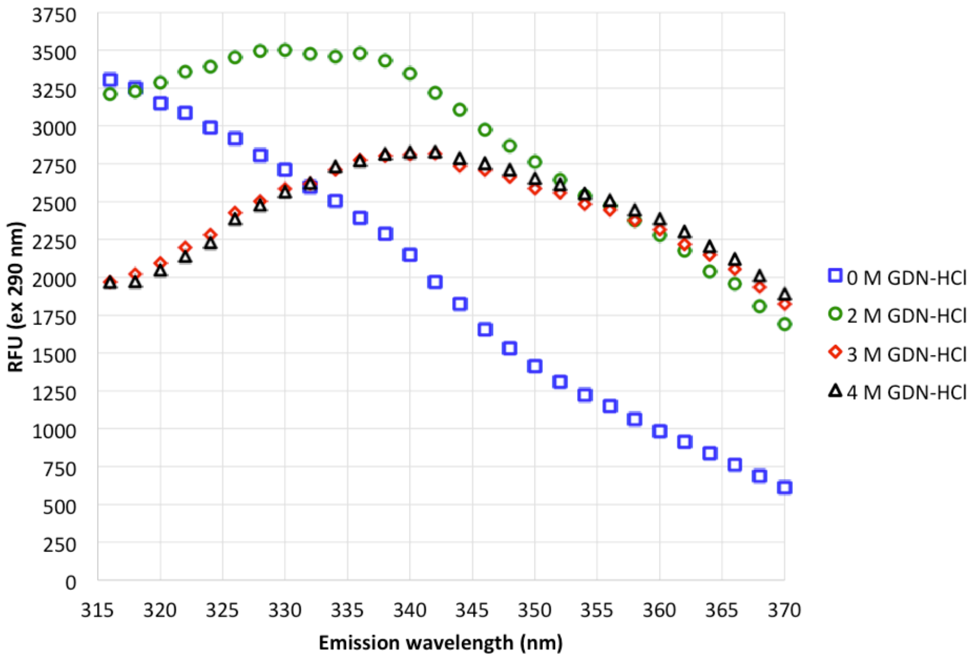


**Fig. S3: AmiE WT tryptophan emission spectra as a function of GDN-HCl concentration**. 52 μM Protein was incubated in PBS with the indicated concentration of GDN-HCl at 4°C for 8 hr before fluorescence measurements. The excitation wavelength was 290 nm, while emission was detected at 315-370 nm. The lack of an isosbestic point between spectra indicates more complicated unfolding than a two-state model.

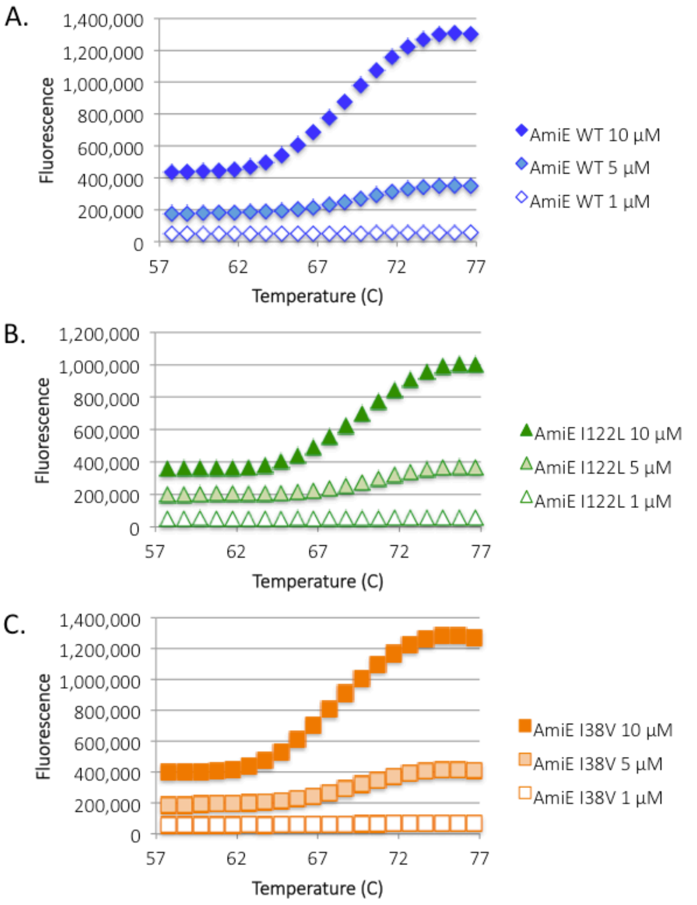


**Fig. S4: Thermal shift analysis of the purified AmiE variants. A.** Melt curves for 10, 5, and 1 μM AmiE WT. **B.** Melt curves for 10, 5, and 1 μM AmiE I122L. **C.** Melt curves for 10, 5, and 1 μM AmiE I38V. All experiments were performed with biological (n=2) and technical (n=3) replicates, and representative curves are shown.

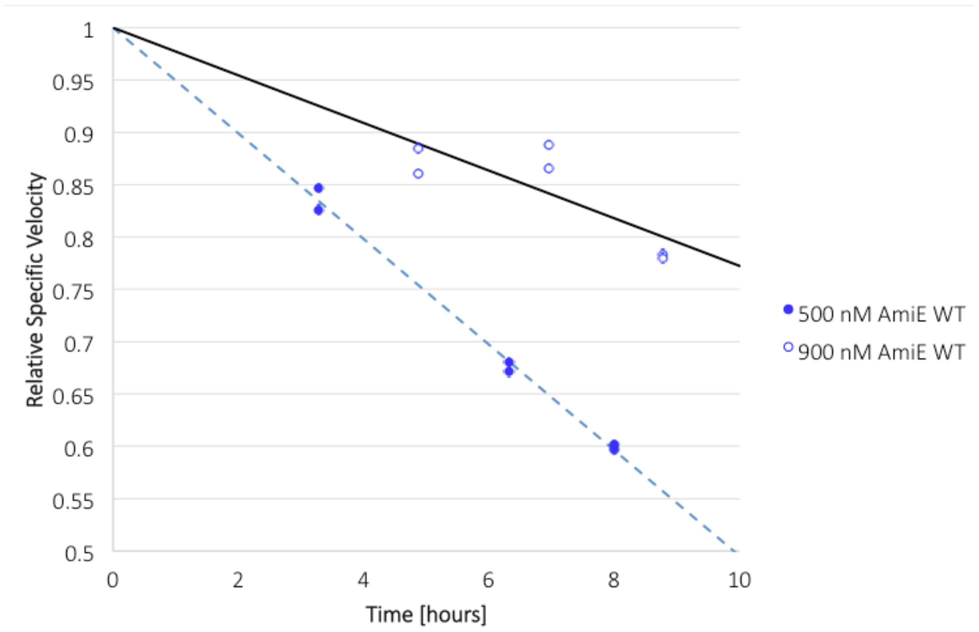


**Fig. S5: Activity loss of AmiE WT following dilution.** Comparison of the relative specific AmiE reaction velocity in a saturating amount of 20 mM acetamide following incubations at dilute concentration in 1x PBS over a time course. Dotted and solid lines represent the best fit linear regressions for the respective data sets (n=6).

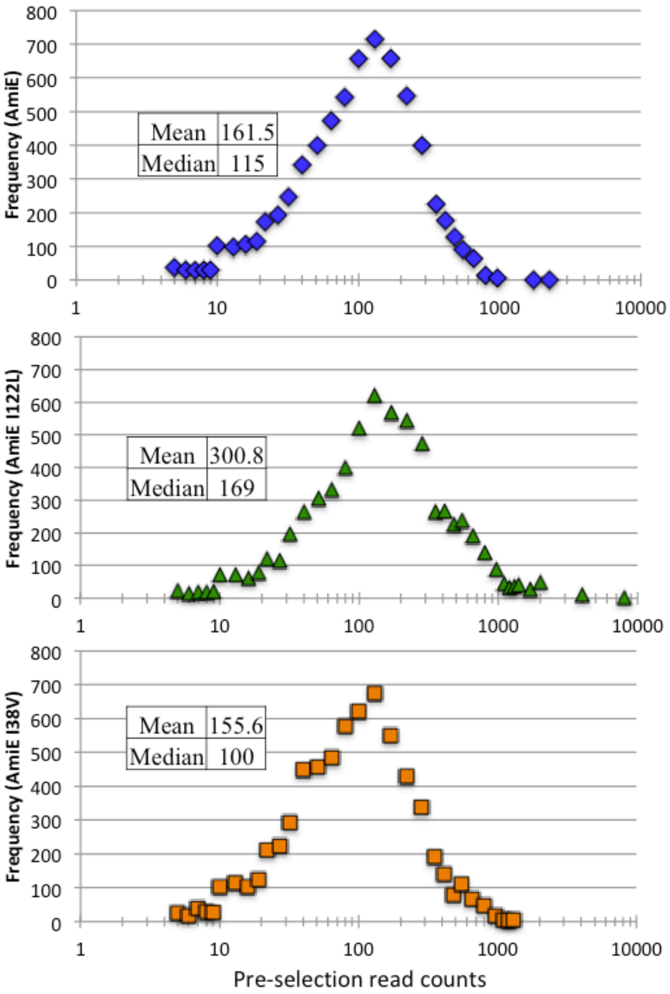


**Fig. S6: Frequency distribution of pre-selection read counts for AmiE libraries.** Mean and median pre-selection read counts for AmiE variants are reported on the plots.

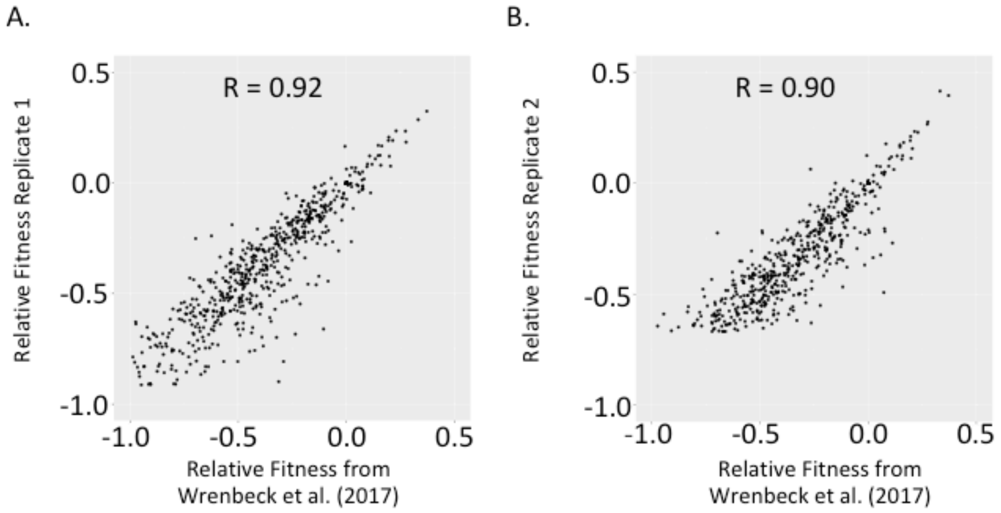


**Fig S7: Deep mutational scanning replicates for AmiE WT compared with previous literature.** Comparison of relative fitness metrics for a AmiE WT mutational library covering residues 171-255 to those from same mutants reported originally in Wrenbeck et al (2017). These replicate deep mutational scans were performed in parallel to AmiE variant growth selections as internal controls. Pearson's correlation coefficients reported on plots. **A**. control for AmiE I122L growth selection. **B**. control for AmiE I38V growth selection.
