## Supporting information part 2 for "Impact of in vivo protein folding probability on local fitness landscapes"

**Corresponding author: Timothy A. Whitehead**

**This PDF file includes:**

Figs. S8 to S14

Tables S1 to S9

**Other supplementary materials for this manuscript include the following:**

Faber_EnzymeStability_ProcessedDatasets.xls

### Supplementary Figures:

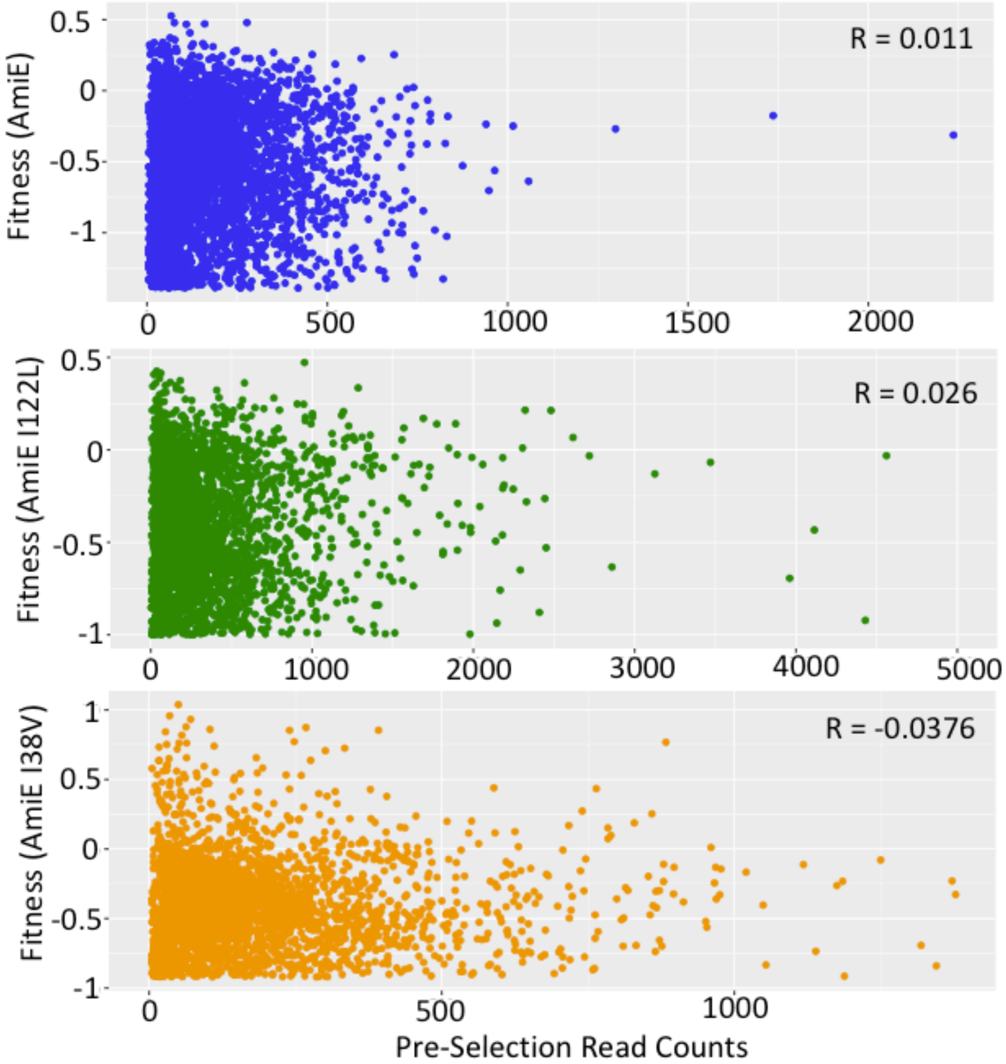

**Fig. S8: Relative fitness metrics for AmiE proteins as a function of pre-selection read counts**. Absolute Pearson's correlation coefficients reported on plots are below 0.04 in all cases, showing that less than 0.2% of the variance can be explained by initial frequency of a given mutant in the library.

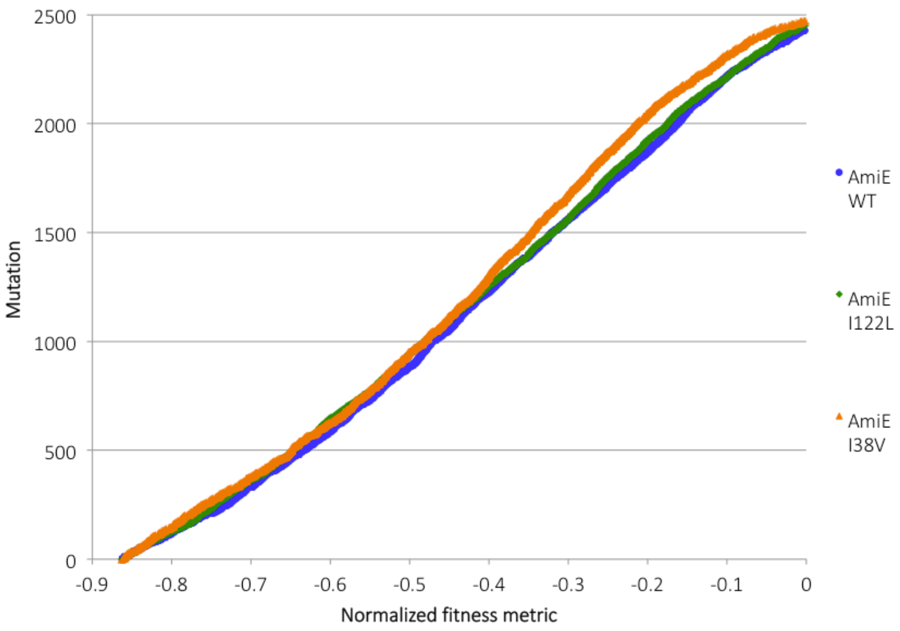

**Fig. S9: Analysis of proportions of deleterious mutations.** Cumulative distribution functions for the deleterious mutations for each enzyme variant are plotted above the lower bounds of experimental measurements.

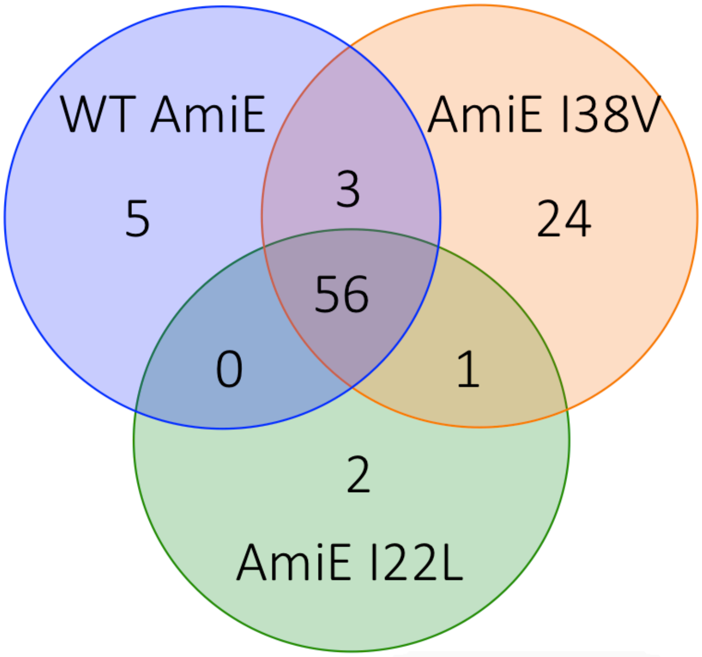

**Fig. S10: Venn diagram of the shared and unique beneficial mutations using the strict cutoff.** Mutations had to improve the growth rate by ≥10% to be classified as a beneficial mutation ($\zeta_i \geq 0.138$).

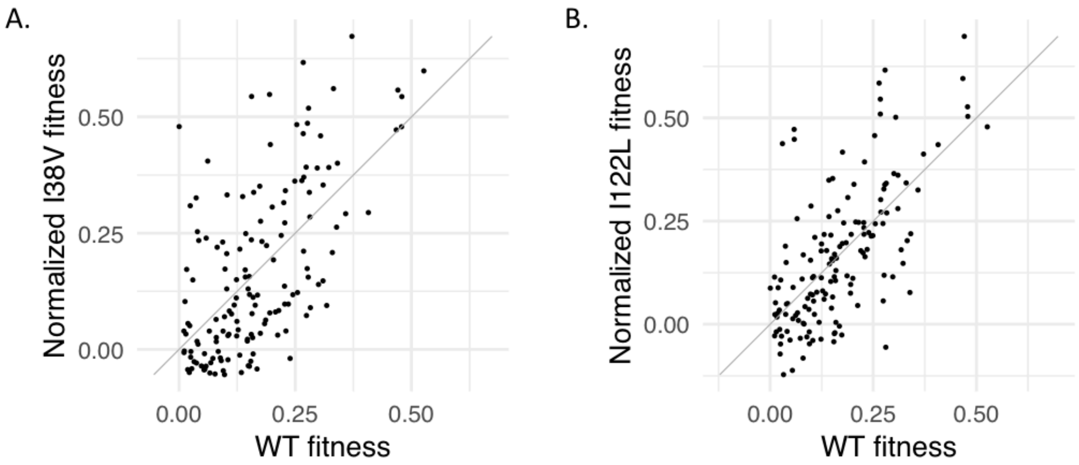

**Fig. S11: Correlation analysis of linear regression normalized shared beneficial mutations.** Comparison of the linear regression normalized fitness metrics for the beneficial mutations shared by all enzymes; solid lines represent the linear function Y = 1X + 0. **A.** Comparison of linear regression normalized AmiE I38V fitness metrics with AmiE WT. **B.** Comparison of linear regression normalized AmiE I122L fitness metrics with AmiE WT.

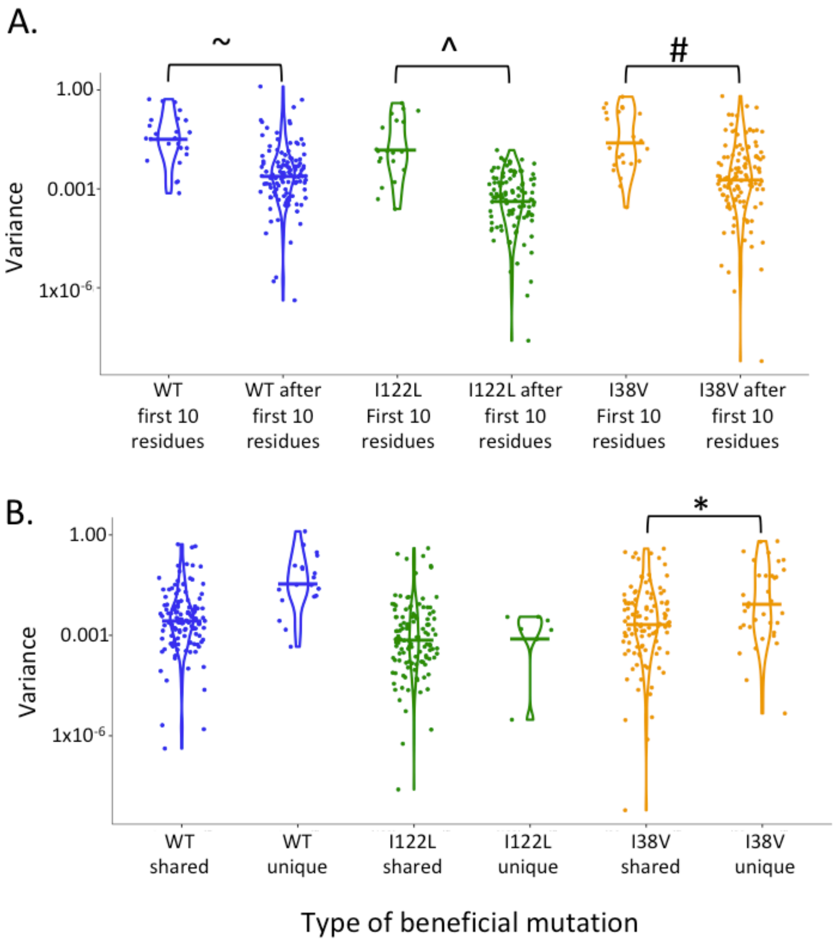

**Fig. S12. Dot plots of fitness effect synonymous codon variances for beneficial mutations. A.** Dot plots of the fitness effect synonymous codon variances for either the beneficial mutations located in the first 10 residues, or all other beneficial mutations after the first 10 residues (95% confidence interval cutoffs). The distribution of the variances is represented by the violin plot overlay and the colored marker lines represent the mean variance for the population, in all three enzymes the first 10 residues have significantly high variance than those outside of the window: ~ = WT p-value = 0.02, ^ = I122L p-value = 0.01, # = I38V p-value = 0.011 from students t-test. **B.** Dot plots of the fitness effect synonymous codon variances for the shared and unique beneficial mutations for the respective enzymes (95% confidence interval cutoffs) The distribution of the variances is represented by the violin plot overlay and the colored marker lines represent the mean variance for the population. I38V has a significantly high variance in the unique beneficial mutations: * = I38V p-value = 0.049 from students t-test.

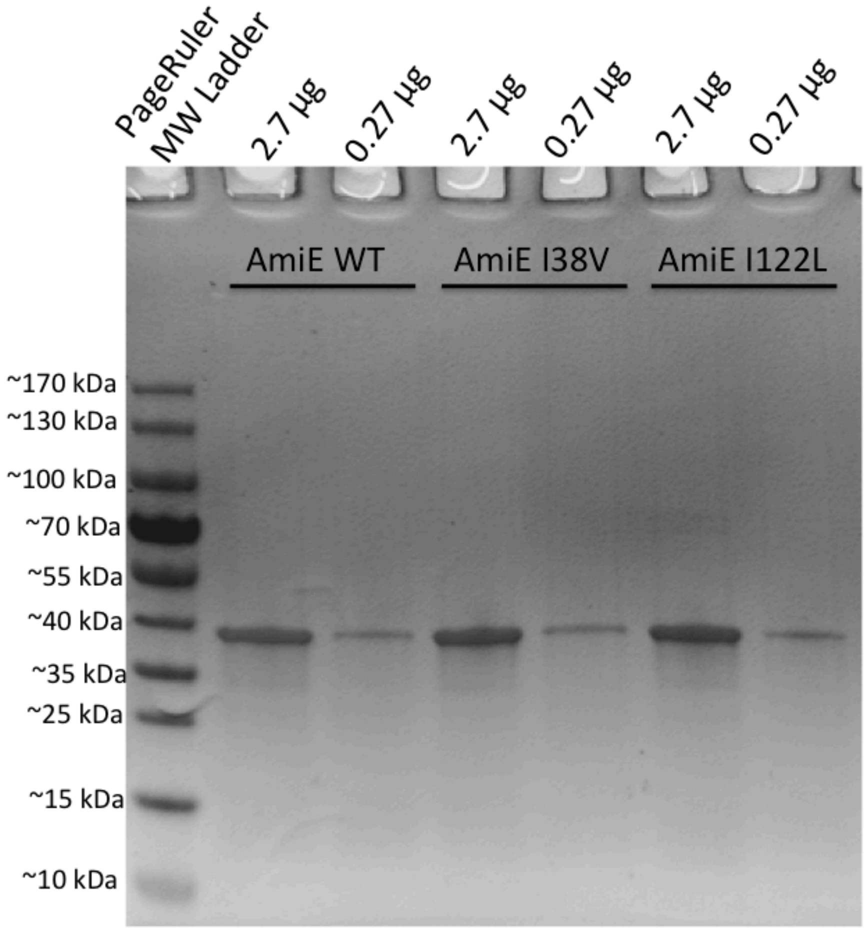

**Fig. S13: SDS-PAGE analysis of the purity of the purified AmiE variants.** Samples were denatured in SDS-PAGE loading buffer (Laemmli's buffer supplemented to 1.5% β-mercaptoethanol at 1x) at 98° for 10 minutes. Samples run on 4-20% Mini-PROTEAN® TGX™ precast gels (Bio-Rad) at 120 V for 1 hour and were washed and stained with SimplyBlue SafeStain (Thermo-Fisher) as described by the manufacturer. Molecular weight ruler used is the PageRuler Prestained Protein Ladder (10-180 kDa, Thermo-Fisher).

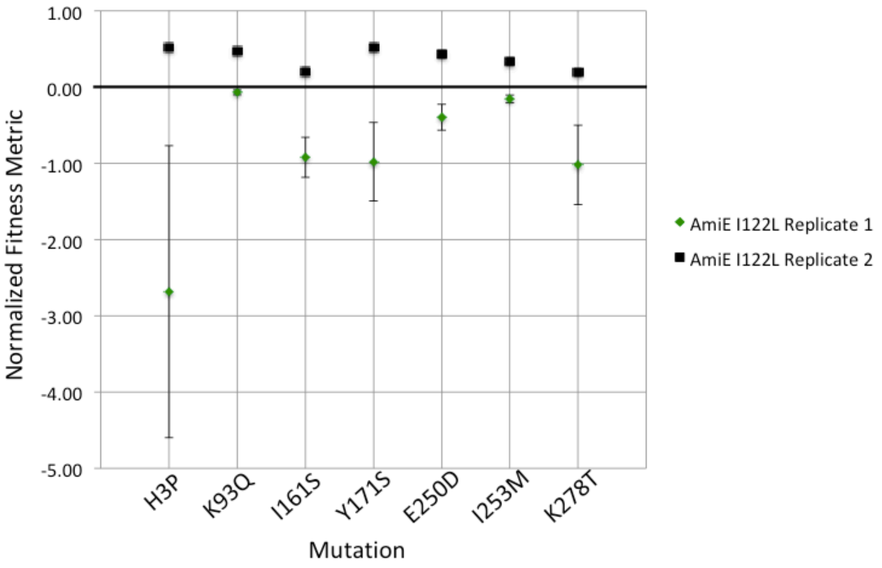

**Fig. S14: Comparison of AmiE I122L outlier mutation technical replicates.** Mutations removed from analysis have their fitness metrics for the respective technical replicates shown, error bars represent the 99.977% confidence intervals (3.5x σ) for the fitness metrics. This indicates that errors such as these should occur less than once per dataset for the size of our experiment.

### Supplementary Tables:

**Table S1: Relative fitness for I38V and I22L synonymous codons in the AmiE WT background.** Green highlight indicates the codons used for the deep mutational scans presented in this work. Data is from Wrenbeck et al (2017).

| | Relative Fitness |
|---|---|
| I38V codon (GTT) | -0.17 |
| I38V codon (GTG) | -0.43 |
| I38V codon (GTC) | -1.17 |
| I38V codon (GTA) | -1.26 |
| I38V average | -0.35 |

| | Relative Fitness |
|---|---|
| I122L codon (TTA) | -0.89 |
| I122L codon (TTG) | -0.69 |
| I122L codon (CTT) | -0.46 |
| I122L codon (CTC) | -0.31 |
| I122L codon (CTA) | -0.54 |
| I122L codon (CTG) | -0.69 |
| I122L average | -0.53 |

**Table S2: Biophysical analysis of the AmiE variants.** Error values reported are 1 s.d. except in the kinetic analysis where 95% confidence interval is given.

| | AmiE Variant | | |
|---|---|---|---|
| | **WT** | **I122L** | **I38V** |
| Rosetta FilterScan score^ | 0.00 | 0.40 | 2.02 |
| Relative $k_{cat}^{*}$ | 1.00 ± 0.06 | 1.08 ± 0.06 | 1.02 ± 0.08 |
| $k_{cat}$ p-value# | 1.00 | 0.57 | 0.79 |
| Relative $K_M^{*}$ | 1.00 ± 0.12 | 0.89 ± 0.15 | 0.82 ± 0.20 |
| $K_M$ p-value# | 1.00 | 0.64 | 0.32 |
| Relative protein yieldŦ | 1 ± 0.06 | 0.26 ± 0.04 | 0.59 ± 0.08 |
| Refolded enzyme velocity normalized to folded enzyme (%) | 14.2 ± 2.47 | 0.114 ± 0.0494 | 0.064 ± 0.0250 |
| Growth Rate in M9 ($hr^{-1}$) - pEDA2 plasmid | 0.78 ± 0.01 | 0.79 ± 0.01 | 0.78 ± 0.02 |
| Growth Rate in Selection Media ($hr^{-1}$) - pEDA2 plasmid | 0.47 ± 0.02 | 0.18 ± 0.01 | 0.11 ± 0.01 |
| Growth Rate in M9 ($hr^{-1}$) - pAG plasmid | 0.79 ± 0.04 | 0.81 ± 0.03 | 0.79 ± 0.01 |
| Growth Rate in Selection Media ($hr^{-1}$) - pAG plasmid | 0.72 ± 0.02 | 0.40 ± 0.03 | 0.33 ± 0.02 |

^ >0 represents a less favorable score to the starting wild-type structure (i.e. decreased likelihood folding)
[*] Relative to AmiE WT

[#] Calculated for variant compared to AmiE WT
[Ŧ] Expressed via Studiers autoinduction

**Table S3: Thermal shift analysis data and statistics.** Summary of thermal melting temperatures ($T_m$) determined using Boltzmann curve fit of a 2-state transition. Students T-Test p-values are reported for tests comparing AmiE WT with the I38V and I122L variants (N = 6 from two biological replicates of each enzyme; ± indicates 1 s.d.).

| Variant | Assaying Concentration (µM) | $T_m$ ($^{o}$C) | p-value | Assaying Concentration (µM) | $T_m$ ($^{o}$C) | p-value | Assaying Concentration (µM) | $T_m$ ($^{o}$C) | p-value |
|---|---|---|---|---|---|---|---|---|---|
| AmiE WT | 1 | N/A | - | 5 | 69.6 ± 0.3 | - | 10 | 68.8 ± 0.2 | - |
| AmiE I122L | 1 | N/A | N/A | 5 | 69.2 ± 1.3 | 0.42 | 10 | 68.7 ± 1 | 0.74 |
| AmiE I38V | 1 | N/A | N/A | 5 | 69.3 ± 0.4 | 0.2 | 10 | 68.5 ± 0.4 | 0.16 |

**Table S4: DNA sequences of transcriptional elements in plasmids used for deep mutational scans.** Differences between sequences are shown in red, the RBS in the 5' UTR is underlined.

| | -35 Promoter | -10 Promoter | 5' UTR |
|---|---|---|---|
| pEDA2 – AmiE | TTCCCG | TAATAT | TCAGGGAGACCACAACGG TTTCCCTCTACAAATAATT TTGTTTAACTTTCTAGAAA TAATTTTGTTTAACTT TAAGAAGTTTTTATACAT |
| pAG – AmiE I38V | TTGGCG | TACAAT | |
| pAG – AmiE I122L | TTGGCG | TACAAT | |

**Table S5: Summary of MG1655 rph+ transformants obtained during selection strain mutant library preparation.** 5 x $10^4$ transformants corresponds to >99% theoretical library coverage.

| Selection Strain SSM Library Preparation | Transformants Obtained |
|---|---|
| AmiE I122L pAG Tile 1 | 15 x $10^5$ |
| AmiE I122L pAG Tile 2 | 19x $10^5$ |
| AmiE I122L pAG Tile 3 | 7.5 x $10^5$ |
| AmiE I122L pAG Tile 4 | 23 x $10^5$ |
| AmiE I38V pAG Tile 1 | 6.9 x $10^5$ |
| AmiE I38V pAG Tile 2 | 5.8 x $10^5$ |
| AmiE I38V pAG Tile 3 | 7.5 x $10^5$ |
| AmiE I38V pAG Tile 4 | 7.8 x $10^5$ |
| WT amiE pEDA2 Tile 3 | 13 x $10^5$ |

**Table S6: Summary of mutational library statistics.**

| | | | | | |
|---|---|---|---|---|---|
| Screen | Growth | | | | |
| Enzyme | AmiE WT | | | | |
| Tile Number | 1 | 2 | 3 | 4 | Cumulative |
| Residues | 1-85 | 86-170 | 171-255 | 256-341 | 1-341 |
| Population | Growth Selected | Growth Selected | Growth Selected | Growth Selected | Growth Selected |
| Number of mutated codons | 85 | 85 | 85 | 86 | 341 |
| Reference sequencing reads post quality filter | 448689 | 393052 | 561875 | 511337 | 1914953 |
| Selected sequencing reads post quality filter | 1454211 | 3016045 | 2101231 | 1691121 | 8262608 |
| **Fold oversampling of codon combinations** | | | | | Average |
| Reference total | 81 | 71.6 | 101.8 | 91.6 | 86.5 |
| Selected total | 256.6 | 544 | 378.1 | 292.1 | 367.7 |
| Reference non-synonymous | 50.8 | 38.3 | 54.5 | 55.8 | 49.9 |
| Selected non-synonymous | 79.8 | 95.7 | 57.4 | 95 | 82.0 |
| **Percent of reads with:** | | | | | Average |
| No nonsynonymous mutations | 37.6 | 46.9 | 46.7 | 39.4 | 42.7 |
| One nonsynonymous mutation | 60.6 | 52.2 | 51.9 | 59.1 | 56.0 |
| Multiple nonsynonymous mutation | 1.8 | 0.9 | 1.4 | 1.4 | 1.4 |
| Coverage of possible single nonsynonymous mutations | 95.2 | 93.9 | 89 | 94.7 | 93.2 |
| Screen | Growth | | | | |
| Enzyme | AmiE I122L | | | | |
| Tile Number | 1 | 2 | 3 | 4 | Cumulative |
| Residues | 1-85 | 86-114, 131, 133-170 | 171-255 | 256-341 | 1-114,131,133-341 |
| Population | Growth Selected | Growth Selected | Growth Selected | Growth Selected | Growth Selected |
| Number of mutated codons | 85 | 68 | 85 | 86 | 324.0 |
| Reference sequencing reads post quality filter | 674718 | 593530 | 897746 | 1214190 | 3380184 |
| Selected sequencing reads post quality filter | 1286349 | 1002404 | 1115963 | 931995 | 4336711 |
| **Fold oversampling of codon combinations** | | | | | Average |
| Reference total | 118.7 | 99 | 146.8 | 210 | 143.6 |
| Selected total | 221.8 | 176.5 | 198.1 | 156 | 188.1 |
| Reference non-synonymous | 76.1 | 61.5 | 90.1 | 137 | 91.2 |
| Selected non-synonymous | 71.6 | 35.8 | 37.8 | 63.6 | 52.2 |
| **Reference population percent of reads with:** | | | | | Average |
| No nonsynonymous mutations | 35.4 | 35.3 | 35.2 | 34 | 35.0 |
| One nonsynonymous mutation | 60.4 | 55.4 | 53.7 | 61.2 | 57.7 |
| Multiple nonsynonymous mutation | 4.3 | 9.3 | 11.1 | 4.8 | 7.4 |
| Coverage of possible single nonsynonymous mutations | 94.5 | 92.5 | 93 | 94.8 | 93.7 |
| Screen | Growth | | | | |
| Enzyme | AmiE I38V | | | | |
| Tile Number | 1 | 2 | 3 | 4 | Cumulative |
| Residues | 1-31, 45-85 | 86-170 | 171-255 | 256-341 | 1-31, 45-341 |
| Population | Growth Selected | Growth Selected | Growth Selected | Growth Selected | Growth Selected |
| Number of mutated codons | 72 | 85 | 85 | 86 | 328 |
| Reference sequencing reads post quality filter | 550823 | 500238 | 525635 | 490442 | 2067138 |
| Selected sequencing reads post quality filter | 1373021 | 1060307 | 991922 | 1129295 | 4554545 |
| **Fold oversampling of codon combinations** | | | | | Average |
| Reference total | 100.7 | 91.5 | 96.1 | 88.6 | 94.2 |
| Selected total | 236.5 | 189 | 173.1 | 185 | 195.9 |
| Reference non-synonymous | 50.3 | 45.7 | 48.3 | 46 | 47.6 |
| Selected non-synonymous | 96.4 | 42.6 | 69.8 | 81 | 72.5 |
| **Percent of reads with:** | | | | | Average |
| No nonsynonymous mutations | 50.5 | 50.5 | 50.3 | 48.6 | 50.0 |
| One nonsynonymous mutation | 48.9 | 48.9 | 49.2 | 50.8 | 49.5 |
| Multiple nonsynonymous mutation | 0.6 | 0.5 | 0.6 | 0.6 | 0.6 |
| Coverage of possible single nonsynonymous mutations | 96.7 | 88.6 | 87.9 | 93.9 | 91.8 |

**Table S7: Goodness-of-fit test statistics for distribution fittings.** For all enzymes the determined p-values indicate a failure to reject the null hypotheses that they can be fit by a general Pareto distribution with a negative shape parameter. All calculations were performed as in Wrenbeck et al. (4).

| General Pareto distribution | AmiE WT | AmiE I122L | AmiE I38V |
|---|---|---|---|
| bootstrap test p-value | 0.16 | 0.29 | 0.10 |
| shape parameter | -0.28 | -0.33 | -0.32 |

**Table S8: Deleterious empirical cumulative distribution function (ECDF) analysis statistics.** A table of the two-sample Kolmogorov-Smirnoff test results for comparing the deleterious mutation ECDFs for the three enzymes.

| Pairing (x-y) | K-S Test | D | p-value | Null hypothesis | Alternative hypothesis | AmiE WT (N) | AmiE variant (N) |
|---|---|---|---|---|---|---|---|
| AmiE WT – AmiE I38V | two sided | 0.058 | 0.0005 | is equal to | not equal | 2428 | 2472 |
| AmiE WT – AmiE I38V | greater | 0.002 | 0.99 | x not greater than y | the CDF of x lies above that of y | 2428 | 2472 |
| AmiE WT – AmiE I38V | less | 0.058 | 0.0002 | x not less than y | the CDF of x lies below that of y | 2428 | 2472 |
| AmiE WT – AmiE I122L | two sided | 0.024 | 0.48 | is equal to | not equal | 2428 | 2450 |
| AmiE WT – AmiE I122L | greater | 0.010 | 0.77 | x not greater than y | the CDF of x lies above that of y | 2428 | 2450 |
| AmiE WT – AmiE I122L | less | 0.024 | 0.24 | x not less than y | the CDF of x lies below that of y | 2428 | 2450 |

**Table S9: Inner and outer primers for PCR reactions for Illumina sequencing.** Red indicates overhang regions for attaching Illumina adapter primers (inner PCR primers) or overhangs for attaching to inner PCR product (outer PCR primers), black is the overlap region in the gene or the barcode, blue is the Illumina adapter.

| **Inner PCR primers** | **Sequence (5' to 3')** |
|---|---|
| Fwd_Tile_1 | gttcagagttctacagtccgacgatcttaactttaagaagtttttatacat |
| Fwd_Tile_2 | gttcagagttctacagtccgacgatcggcgaagaaacggaa |
| Fwd_Tile_3 | gttcagagttctacagtccgacgatcctgcgatgacggtaat |
| Fwd_Tile_4 | gttcagagttctacagtccgacgatcaagaaatgggcattcaatac |
| Rev_Tile_1 | ccttggcacccgagaattccaaagcacggctaaagat |
| Rev_Tile_2 | ccttggcacccgagaattccactctccaaatttccggata |
| Rev_Tile_3 | ccttggcacccgagaattccacagagacaactgcgc |
| Rev_Tile_4 | ccttggcacccgagaattccatggtggtgctcgag |
| **Illumina outer primer adapter** | aatgatacggcgaccaccgagatctacacgttcagagttctacagtccga |
| **Illumina outer PCR adapters and barcodes** | |
| RPI37 (AmiE WT unselected, Tile 1) | caagcagaagacggcatacgagatATTCCGgtgactggagttccttggcacccgagaattcca |
| RPI22 (AmiE WT unselected, Tile 2) | caagcagaagacggcatacgagatCGTACGgtgactggagttccttggcacccgagaattcca |
| RPI39 (AmiE WT unselected, Tile 3) | caagcagaagacggcatacgagatGTATAGgtgactggagttccttggcacccgagaattcca |
| RPI40 (AmiE WT unselected, Tile 4) | caagcagaagacggcatacgagatTCTGAGgtgactggagttccttggcacccgagaattcca |
| RPI41 (AmiE WT selected, Tile 1 replicate 1) | caagcagaagacggcatacgagatGTCGTCgtgactggagttccttggcacccgagaattcca |
| RPI38 (AmiE WT selected, Tile 1 replicate 2) | caagcagaagacggcatacgagatAGCTAGgtgactggagttccttggcacccgagaattcca |
| RPI33 (AmiE WT selected, Tile 2 replicate 1) | caagcagaagacggcatacgagatCGCCTGgtgactggagttccttggcacccgagaattcca |
| RPI34 (AmiE WT selected, Tile 2 replicate 2) | caagcagaagacggcatacgagatGCCATGgtgactggagttccttggcacccgagaattcca |
| RPI43 (AmiE WT selected, Tile 3 replicate 1) | caagcagaagacggcatacgagatGCTGTAgtgactggagttccttggcacccgagaattcca |
| RPI40 (AmiE WT selected, Tile 3 replicate 2) | caagcagaagacggcatacgagatTCTGAGgtgactggagttccttggcacccgagaattcca |
| RPI44 (AmiE WT selected, Tile 4 replicate 1) | caagcagaagacggcatacgagatATTATAgtgactggagttccttggcacccgagaattcca |
| RPI41 (AmiE WT selected, Tile 4 replicate 2) | caagcagaagacggcatacgagatGTCGTCgtgactggagttccttggcacccgagaattcca |
| RPI20 (AmiE I38V unselected, Tile 1) | caagcagaagacggcatacgagatGGCCACgtgactggagttccttggcacccgagaattcca |
| RPI21 (AmiE I38V unselected, Tile 2) | caagcagaagacggcatacgagatCGAAACgtgactggagttccttggcacccgagaattcca |
| RPI27 (AmiE I38V unselected, Tile 3) | caagcagaagacggcatacgagatAGGAATgtgactggagttccttggcacccgagaattcca |
| RPI34 (AmiE I38V unselected, Tile 4) | caagcagaagacggcatacgagatGCCATGgtgactggagttccttggcacccgagaattcca |
| RPI37 (AmiE I38V selected, Tile 1 replicate 1) | caagcagaagacggcatacgagatATTCCGgtgactggagttccttggcacccgagaattcca |
| RPI38 (AmiE I38V selected, Tile 1 replicate 2) | caagcagaagacggcatacgagatAGCTAGgtgactggagttccttggcacccgagaattcca |
| RPI16 (AmiE I38V selected, Tile 2 replicate 1) | caagcagaagacggcatacgagatGGACGGgtgactggagttccttggcacccgagaattcca |
| RPI14 (AmiE I38V selected, Tile 2 | caagcagaagacggcatacgagatGGAACTgtgactggagttccttggcacccgagaattcca |

| | |
|---|---|
| replicate 2) | |
| RPI48 (AmiE I38V selected, Tile 3 replicate 1) | caagcagaagacggcatacgagatTGCCGAgtgactggagttccttggcacccgagaattcca |
| RPI23 (AmiE I38V selected, Tile 3 replicate 2 ) | caagcagaagacggcatacgagatCCACTCgtgactggagttccttggcacccgagaattcca |
| RPI32 (AmiE I38V selected, Tile 4 replicate 1) | caagcagaagacggcatacgagatTGAGTGgtgactggagttccttggcacccgagaattcca |
| RPI44 (AmiE I38V selected, Tile 4 replicate 2) | caagcagaagacggcatacgagatATTATAgtgactggagttccttggcacccgagaattcca |
| RPI19 (AmiE WT control, AmiE I38V selection, unselected, Tile 3) | caagcagaagacggcatacgagatTTTCACgtgactggagttccttggcacccgagaattcca |
| RPI26 (AmiE WT control, AmiE I38V selection, selected, Tile 3) | caagcagaagacggcatacgagatGCTCATgtgactggagttccttggcacccgagaattcca |
| RPI33 (AmiE I122L unselected, Tile 1, sequencing runs 1 and 2) | caagcagaagacggcatacgagatCGCCTGgtgactggagttccttggcacccgagaattcca |
| RPI46 (AmiE I122L unselected, Tile 2, sequencing runs 1 and 2) | caagcagaagacggcatacgagatTCGGGAgtgactggagttccttggcacccgagaattcca |
| RPI40 (AmiE I122L unselected, Tile 3, sequencing runs 1and 2) | caagcagaagacggcatacgagatTCTGAGgtgactggagttccttggcacccgagaattcca |
| RPI29 (AmiE I122L unselected, Tile 4, sequencing runs 1 and 2) | caagcagaagacggcatacgagatTAGTTGgtgactggagttccttggcacccgagaattcca |
| RPI17 (AmiE I122L selected, Tile 1 replicate 1) | caagcagaagacggcatacgagatCTCTACgtgactggagttccttggcacccgagaattcca |
| RPI32 (AmiE I122L selected, Tile 1 replicate 2) | caagcagaagacggcatacgagatTGAGTGgtgactggagttccttggcacccgagaattcca |
| RPI12 (AmiE I122L selected, Tile 2 replicate 1) | caagcagaagacggcatacgagatTACAAGgtgactggagttccttggcacccgagaattcca |
| RPI42 (AmiE I122L selected, Tile 2 replicate 2) | caagcagaagacggcatacgagatCGATTAgtgactggagttccttggcacccgagaattcca |
| RPI25 (AmiE I122L selected, Tile 3 replicate 1) | caagcagaagacggcatacgagatATCAGTgtgactggagttccttggcacccgagaattcca |
| RPI46 (AmiE I122L selected, Tile 3 replicate 2 ) | caagcagaagacggcatacgagatTCGGGAgtgactggagttccttggcacccgagaattcca |
| RPI31 (AmiE I122L selected, Tile 4 replicate 1) | caagcagaagacggcatacgagatATCGTGgtgactggagttccttggcacccgagaattcca |
| RPI15 (AmiE I122L selected, Tile 4 replicate 2) | caagcagaagacggcatacgagatTGACATgtgactggagttccttggcacccgagaattcca |
| RPI7 (AmiE WT control, AmiE I122L selection, unselected, Tile 3) | caagcagaagacggcatacgagatGATCTGgtgactggagttccttggcacccgagaattcca |
| RPI10 (AmiE WT control, AmiE I122L selection, selected, Tile 3) | caagcagaagacggcatacgagatAAGCTAgtgactggagttccttggcacccgagaattcca |

**Additional data** (Faber_EnzymeStability_ProcessedDatasets.xls)

Contains spreadsheets with the processed data for this paper, it includes: fitness metric heatmaps, analysis of shared and unique beneficial mutations, analysis of mutations with codon fitness metric disparities.
